# Unveiling the Multifaceted Networks of the TMS-Targeted Left Prefrontal Cortex for Precision Neurostimulation

**DOI:** 10.1101/2025.11.19.689337

**Authors:** Lya K. Paas Oliveros, Timm B. Poeppl, Niels Reuter, Kaustubh R. Patil, Sarah Kreuzer, Nga Yan Tse, Robin F. H. Cash, Felix Hoffstaedter, Simon B. Eickhoff, Veronika I. Müller

## Abstract

The left dorsolateral prefrontal cortex (lDLPFC) is the standard transcranial magnetic stimulation (TMS) target for treatment-resistant depression (TRD), yet non-response rates remain high. TMS efficacy has been linked to the stimulation site’s functional connectivity, particularly its anti-correlation with the subgenual cingulate cortex (SGC). While this pragmatic strategy has demonstrated clinical utility, it offers limited insight into the functional organization of the lDLPFC, potentially contributing to site-dependent variability in treatment response. Here, we applied connectivity-based parcellation to a region of interest encompassing common TMS targets in the left prefrontal cortex (TMS-PFC). This yielded a TMS-PFC functional atlas in a healthy population, with each subregion characterized by its large-scale networks and behavioral associations. Our results revealed a hierarchical organization: A coarse two-pole antagonism separating anterior-central from superior-posterior subregions, while a finer nine-cluster solution revealed heterogeneity along anterior-posterior and ventral-dorsal axes. Anterior-central areas were strongly anti-correlated with SGC and default-mode network, positively connected with salience, dorsal attention, and control networks, and associated with cognitive control. In contrast, superior-posterior subregions displayed the inverse pattern, while ventral clusters engaged somatomotor and visual networks, and language-related processes. Central and superior-anterior clusters showed associations with inhibition, social cognition, and perceptual functions. Integrating across granularities, we derived a map highlighting the anterior-central subregion as a promising candidate for subsequent targeting studies given its connectivity and behavioral profile, while neighboring subregions showed distinct functions. These findings underscore the hierarchical and heterogeneous organization of TMS-PFC and provide a network-informed atlas for future research on individualized, symptom-specific TMS targeting.

**Highlights:**

- TMS targets within the left PFC encompass nine functionally distinct subregions
- Subregions differ in SGC, DMN, salience, attention, and control network connectivity
- Anterior-central and superior-posterior subregions show antagonistic profiles
- The anterior-central subregion shows the strongest SGC-anticorrelated connectivity
- Findings provide a network-informed reference for potential TMS targeting

## 1 Introduction

With a lifetime prevalence of up to 20%, depression remains a leading cause of disability worldwide, carrying substantial personal, societal, and economic burdens (Institute for Health Metrics and Evaluation, 2020; World Health Organization, 2017). These burdens are amplified in treatment-resistant depression (TRD), typically defined as inadequate response to at least two antidepressant treatments (European Medicines Agency, 2025; U.S. Food and Drug Administration, 2018). Transcranial magnetic stimulation (TMS) targeting the left dorsolateral prefrontal cortex (lDLPFC) has become a standard treatment to ameliorate TRD (Cash, Weigand, et al., 2021; Fitzgerald et al., 2016). Despite its clinical efficacy, response and remission rates remain variable, with approximately 50% of patients showing no meaningful improvement (Blumberger et al., 2018; Fitzgerald et al., 2016).

This variability has sparked efforts to optimize the stimulation-site selection. Conventional lDLPFC localization relies on anatomical landmarks or scalp-based coordinates, such as the 5.5-cm method anterior to the motor hotspot or the F3 electrode position (Beam F3) (for a detailed review, see Cash, Weigand, et al., 2021). Although these scalp-based heuristics have undergone incremental refinement to improve anatomical accuracy, this has not translated into better treatment outcomes (Sakreida et al., 2025). Part of this challenge is that the lDLPFC itself does not constitute a single, uniformly defined anatomical entity, but rather a heterogeneous region whose boundaries and subdivisions vary across cytoarchitectonic, functional, and stimulation-oriented frameworks (Brodmann, 1909; Bruno et al., 2024; Cash, Weigand, et al., 2021; Cieslik et al., 2013; Li et al., 2024; Petrides & Pandya, 1999; Sallet et al., 2013). Indeed, as neuronavigation systems have been introduced, a wide spectrum of left prefrontal TMS targets has emerged, extending from superior-posterior to anterior-inferior sections beyond the classical lDLPFC (Cash, Weigand, et al., 2021). This is further compounded by the marked inter-individual variability in DLPFC’s morphology, neural function, and structural and functional connectivity (Cieslik et al., 2013; Doucet et al., 2019; M. D. Fox et al., 2013; Friedman & Miyake, 2017; Mueller et al., 2013; Panikratova et al., 2020; Rajkowska & Goldman-Rakic, 1995; Sallet et al., 2013).

Cytoarchitectonic work has long subdivided the DLPFC into BA9 and BA46 (Brodmann, 1909; Sarkissov et al., 1955; von Economo & Koskinas, 1925) and further into dorsal and ventral transition zones (BA9/46d and BA9/46v) (Petrides & Pandya, 1999), with a more recent map distinguishing nine microstructurally subregions (Bruno et al., 2024). Functional parcellation efforts add further complexity, identifying from two to ten DLPFC subregions with distinct connectivity profiles (Cieslik et al., 2013; Jung et al., 2022; Li et al., 2024; Sallet et al., 2013). However, each of these studies suffers from some methodological limitation, such as small sample sizes (Sallet et al., 2013), non-adult cohorts (Li et al., 2024), right-hemisphere focus (Cieslik et al., 2013; Sallet et al., 2013) or spherical representations (Jung et al., 2022). The spatial extent and definition of the DLPFC used in these studies also vary substantially, from narrowly defined right-hemispheric areas (Cieslik et al., 2013) to large sections of the frontal cortex (Sallet et al., 2013). Thus, despite these multiple parcellation efforts, none has systematically parcellated and characterized the left prefrontal area encompassing the full range of clinically used TMS-target sites.

While none of the parcellation efforts have been translated to TMS-site selection, connectivity-guided targeting strategies have more recently gained popularity, aiming to personalize stimulation sites using intrinsic functional connectivity (Cash et al., 2019; Cole et al., 2022; M. D. Fox et al., 2012, 2013; Weigand et al., 2018). The prevailing approach identifies the lDLPFC site with the strongest negative resting-state functional connectivity (RSFC) to the subgenual cingulate cortex (SGC) (Cash et al., 2019; Cole et al., 2022; M. D. Fox et al., 2012, 2013; Weigand et al., 2018). This approach draws on circuit-level accounts of cortico-limbic dysregulation, in which SGC has been highlighted as a key region in the maintenance and development of TRD pathophysiology (Fitzgerald et al., 2008; Koenigs & Grafman, 2009; Mayberg et al., 2005). SGC hyperactivity and lDLPFC hypoactivity are well-documented, treatment-responsive alterations (Drevets et al., 2008; Koenigs & Grafman, 2009; Mayberg et al., 2005).

However, while the SGC-centric approach has demonstrated promise and has strong neurobiological underpinnings, a more in-depth understanding of the functional architecture of TMS-targeted prefrontal areas and associated behavioral profiles could provide a new foundation for refining precision TMS targeting strategies. Indeed, depression has been consistently associated with aberrant connectivity across long-range brain networks that support affective, cognitive, and somatic domains, including the default mode (DMN), frontoparietal (FPN), dorsal attention (DAN), and salience (SAN) networks (Chai et al., 2023; Prompiengchai & Dunlop, 2025). Importantly, these alterations are not static, and effective treatments, including TMS, can modulate connectivity both locally and distally across distributed systems in ways that track clinical improvement (Beynel et al., 2020; Chai et al., 2023; Drysdale et al., 2017; Prompiengchai & Dunlop, 2025). For instance, reduced connectivity between the stimulated DLPFC and SAN, alongside increased SAN activity (fractional amplitude of low-frequency fluctuations, fALFF) and SAN-posterior DMN (pDMN) connectivity, have been associated with antidepressant response (Godfrey et al., 2022), suggesting that TMS may restore balance within the DMN-FPN-SAN canonical triple network (Godfrey et al., 2022; B. Menon, 2019; V. Menon, 2011). Complementary work has also reported TMS-induced decreases in intra-DMN and DMN–FPN connectivity, alongside increased DMN-limbic coupling (Chai et al., 2023), underscoring the widespread and distributed consequences of stimulation.

Nevertheless, treatment response still remains highly variable, potentially reflecting both inter-individual heterogeneity in network alterations and the symptomatic diversity of depression. Symptom-specific circuit models highlight that different depressive symptoms may rely on dissociable brain circuits (Cash et al., 2023; Siddiqi et al., 2020): Dysphoric symptoms (e.g., sadness, anhedonia, and suicidal ideation) appear to respond better to stimulation of anterolateral DLPFC areas anticorrelated with SGC, whereas anxiety and somatic complaints (e.g., insomnia, irritability, and decreased libido) respond more favorably to stimulation of posterior DLPFC sites functionally connected with mPFC (Siddiqi et al., 2020). Circuits and targets broadly relating to affective versus cognitive symptoms have also been proposed (Cash et al., 2023). Although some of these approaches acknowledge the broader network dysfunction in TRD and the distributed TMS effects via distal circuits, these insights are not yet directly incorporated into targeting methodologies (Cash & Zalesky, 2024; Castrillon et al., 2020; Gratton et al., 2013; Siddiqi et al., 2021).

Together, this literature points to two related gaps. First, targeting methodologies to date provide only a limited picture of the highly heterogeneous functional architecture of the selected prefrontal target region itself, in relation to its subregional boundaries, organizational structure, and integration with large-scale networks. Second, despite evidence that distinct networks and circuits underlie different depressive symptoms, a clinically-anchored functional map of the TMS targets is still lacking. That is, no existing framework systematically maps the intrinsic connectivity profiles of TMS-targeted left prefrontal subregions and situates them within large-scale networks, a step needed not only to better understand the distributed effects of stimulation but also to advance network-informed stimulation strategies that go beyond SGC-centric models. These observations highlight the need for more probabilistic targeting models that capture the complex macrostructural organization of the TMS-targeted left prefrontal cortex (TMS-PFC) and account for the substantial inter-individual variability in its intrinsic connectivity.

Here, we sought to delineate and characterize the functional subregions encompassed by clinically used TMS targets within the left prefrontal cortex. To this end, we defined a region of interest capturing the heterogeneity of stimulation coordinates used in prior studies and applied regional connectivity-based parcellation (rCBP) to identify functionally distinct TMS-PFC subregions from whole-brain resting-state connectivity. Rather than a general-purpose parcellation of the lDLPFC, we aimed for a functional subdivision specifically tailored for neuromodulation applications, determining which intrinsic functional regions underlie clinically used stimulation sites. Each resulting subregion was subsequently characterized by its functional connectivity with SGC and large-scale brain networks, as well as by its associated behavioral domains through quantitative functional decoding. This followed the particular interest of establishing which broader functional systems are reached by clinically used stimulation sites and how these subregions differ in their network embedding and behavioral relevance. This framework provides a robust group-level map of TMS-PFC functional subunits and a foundation for more precise, network-informed TMS targeting strategies.

## 2 Methods

### 2.1 Region of Interest

The region of interest (ROI) was defined around commonly used prefrontal TMS target coordinates in depression (see **Supplementary Methods, Table S1**) (Cash, Cocchi, Lv, Fitzgerald, et al., 2021; Cash, Weigand, et al., 2021; M. D. Fox et al., 2012). The TMS coordinates were mainly located in the middle frontal gyrus and frontal pole, according to the Harvard-Oxford Structural Probability Atlas (Desikan et al., 2006). We computed 2000 (*k-*)nearest neighbors (KNN) around the TMS coordinates, restricted by a gray-matter (GM) mask in MNI152NLin6Asym space with 2 mm isotropic voxels. This value was determined empirically by testing a range of KNN values (1000–6000, in increments of 1000): Smaller KNNs resulted in gaps within the ROI, whereas larger values extended the ROI into superior, ventral, and medial frontal areas beyond the intended TMS-targeted area. To avoid inclusion of primary motor and premotor regions, we excluded the left precentral gyrus from the Harvard-Oxford Structural Probability Atlas (Desikan et al., 2006) and the left dorsal premotor cortex (BA6) from the Jülich-Brain Cytoarchitectonic Atlas (Amunts et al., 2020), based on the maps provided by FSLeyes (v1.4.6) and thresholded at 10% probability. Finally, we median-filtered the binarized ROI mask after the exclusion to reduce isolated voxels and irregular border artifacts by voxel-wise masking. The resulting TMS-PFC seed comprised 4232 voxels (33856 mm^3^; **Figure 1A**) extending mainly over the middle frontal gyrus and frontal pole but also covering small sections of the inferior and superior frontal gyri, according to the Harvard-Oxford Structural Probability Atlas (Desikan et al., 2006).

**Figure 1.**
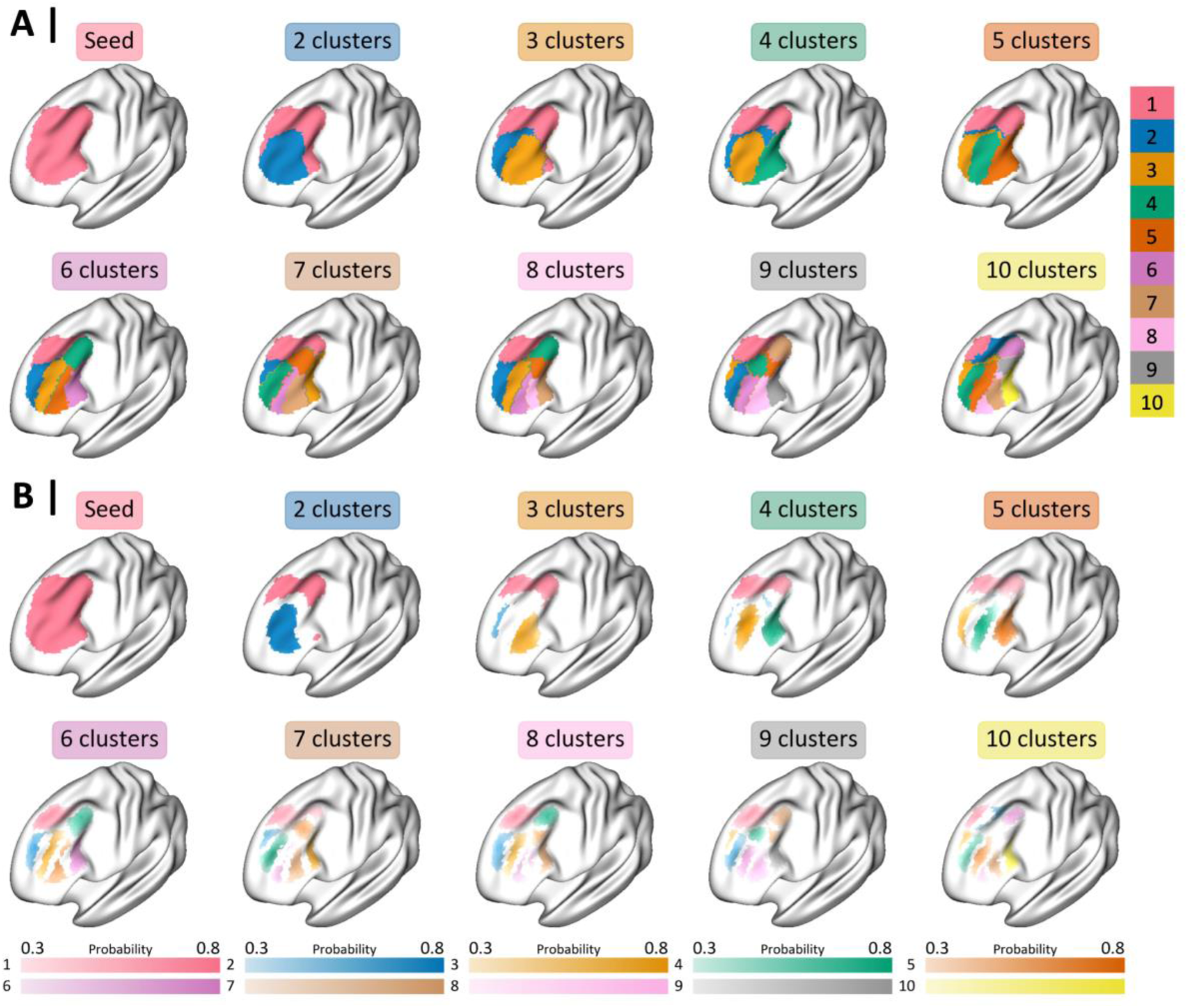
Regional connectivity-based parcellation of the TMS-targeted left prefrontal cortex. **A.** Seed of TMS-PFC and parcellation results into nine cluster solutions (*k* = 2–10) based on RSFC. **B.** Probabilistic maps of each cluster across all parcellation granularities. Maps represent the voxel-wise likelihood of cluster assignment across all subjects after relabeling individual cluster labels with the reference clustering image. Voxels with a probability lower than 30% or those shared between clusters (i.e., spatial overlap) were excluded and labeled with 0 to retain only voxels with sufficient consistency, improve clarity, and reduce ambiguity in cluster boundaries by avoiding overlapping probabilities between clusters in single voxels. **Abbreviations.** RSFC: resting-state functional connectivity, TMS: transcranial magnetic stimulation.

### 2.2 Dataset

Data were obtained from the openly available Human Connectome Project Young Adult (HCP-YA) S1200 release dataset (Van Essen et al., 2013). We included only unrelated participants to avoid heritability effects while maintaining an equal sex distribution. In addition, only participants with all four runs of rs-fMRI, no quality control issues, and diffusion-tensor imaging (DTI) data were included, resulting in a sample of 391 participants (189 female [48.3%], mean age of 28.7 ± 3.8 years [range: 22– 36]). Restriction to subjects with all four rs-fMRI and DTI sessions was due to a parallel project not included in this publication. All participants provided written informed consent, and the reanalysis of the data was approved by the ethics committee of the Medical Faculty at the Heinrich Heine University Düsseldorf (4039 and 2018-317-RetroDEuA).

All HCP scans were acquired on a 3T Siemens “Connectome Skyra” MRI scanner (Siemens AG, Erlangen, Germany) with a 32-channel head coil (Glasser et al., 2013). Structural T1-weighted MRI images were obtained using a 3D multi-echo MPRAGE sequence (TR = 2400 ms, TE = 2.14 ms, TI = 1000 ms, FoV = 224 × 224 mm, flip angle = 8°, 0.7 mm isotropic voxels, 256 slices). The rs-fMRI images were acquired using a whole-brain multi-band gradient-echo echo-planar imaging (TR = 720 ms, TE = 33.10 ms, FoV = 208 × 180 mm, flip angle = 52°, 2 mm isotropic voxels, 72 slices per volume). Each rs-fMRI session acquired 1200 volumes each (14:33 min, eyes open and fixated on a cross-hair) across two consecutive days, with two sessions on each day using opposite phase-encoding directions (right to left and left to right). For this study, we used only the two sessions from day one, as we have planned to use the day-two rs-fMRI sessions for a consecutive project investigating intra-individual reliability.

### 2.3 MRI Preprocessing

We used the publicly available ICA-FIX-denoised rs-fMRI data from HCP-YA, which was preprocessed as described in Glasser et al. (2013). In brief, the raw functional images from each session underwent minimal preprocessing, including corrections for gradient nonlinearity, head motion, and B_0_ distortions. This was followed by registration to the participant’s structural T1-weighted image, spatial normalization into MNI standard space, intensity normalization, and brain masking (Griffanti et al., 2014; Salimi-Khorshidi et al., 2014; Smith et al., 2013). Additionally, independent component analysis (ICA) was combined with the automated component classifier FMRIB’s ICA-based X-noiseifier (FIX) for artifact removal (Beckmann & Smith, 2004; Salimi-Khorshidi et al., 2014). We further processed the data as follows: We regressed out the global mean time courses of white matter and cerebrospinal fluid, as this has been shown to yield highly stable RSFC parcellations (Eickhoff et al., 2015, 2018; Reuter et al., 2020), applied band-pass filtering between 0.01 and 0.08 Hz, and performed spatial smoothing with a 5 mm full-width at half-maximum isotropic Gaussian kernel.

### 2.4 Regional Connectivity-Based Parcellation

To identify functional subregions of the TMS-PFC based on resting-state functional connectivity (RSFC), we applied regional connectivity-based parcellation (rCBP) using published *CBPtools* (Reuter et al., 2020), which was adapted for Python v3.11.8 and our computing environment. Consistent with previous rCBP studies (Genon et al., 2017; Liu et al., 2021; Plachti et al., 2019), this approach groups the voxels within the ROI into distinct subregions (i.e., clusters) through a clustering algorithm based on the similarity of their RSFC patterns. The procedure is performed in two steps: The primary parcellation is computed by-subject and is followed by a group-level parcellation to identify common subregions across participants.

First, the ROI mask was binarized and removed from the target GM mask, which, in turn, was binarized and subsampled, resulting in a target GM mask of 19041 voxels. At the individual level, for each session and voxel of the ROI, Pearson correlation coefficients were computed between the preprocessed voxel time series of the seed and all target voxels, followed by Fisher’s *z*-transformation to derive the RSFC estimates for each session. The individual voxel-level seed-to-target connectivity matrices used for clustering were obtained by averaging the RSFC matrices across both sessions.

For each subject, *k*-means clustering was performed with 256 initializations (Nanetti et al., 2009) and a maximum of 10000 iterations to examine nine levels of granularity (*k* = 2–10; upper limit constrained by computation demands). *K*-means clustering was selected as the individual-level parcellation algorithm, consistent with its widespread use in CBP literature (Genon et al., 2017; Liu et al., 2020; Plachti et al., 2019; Reuter et al., 2020). While alternative algorithms such as spectral and agglomerative clustering are available, their suitability depends on the nature of the data. In particular, spectral clustering, although offering high reproducibility, may impose a strong spatial structure on the data that is not naturally present and is therefore less well-suited for high-level associative brain regions where clusters are not expected to separate cleanly (Eickhoff et al., 2015). Furthermore, no single algorithm has been shown to optimally address all parcellation challenges simultaneously (Arslan et al., 2018), supporting the use of the well-validated *k*-means approach as a reasonable and consistent methodological choice.

To obtain a reference clustering per *k* across subjects, the individual-level cluster labels, i.e., one cluster label per seed voxel per subject, were entered into hierarchical clustering with complete linkage and Hamming distance. Hamming distance was chosen because it is invariant to the arbitrary assignment of cluster labels at the individual level, measuring only the proportion of voxels for which two subjects’ assignments differ (Nguyen & Caruana, 2007; Reuter et al., 2020). Pairwise Hamming distance was computed across all subjects’ label vectors, hierarchical clustering was performed on the resulting distance matrix with complete linkage, and the dendrogram was cut at the requested number of clusters (*k*), yielding a reference clustering for each granularity. This reference clustering was then used to relabel the individual clustering assignments through a permutation approach: For each subject, all possible permutations of cluster label swaps were evaluated against the reference clustering, and the permutation maximizing label agreement with the reference labels was retained to reassign that subject’s cluster labels, ensuring comparability across subjects. The final group-specific parcellation was derived by assigning each voxel the most frequent label across relabeled individual solutions, determined by the mode.

To create a probabilistic map of the individual parcellations, we used the relabeled clustering matrices and computed, for each cluster and each granularity level, the frequency with which each voxel was assigned to the cluster across participants. This yielded voxel-wise probability values reflecting the consistency of cluster assignment at the group level. Brain visualizations were created using Surfplot (Gale et al., 2021; Vos de Wael et al., 2020).

### 2.5 Evaluation of Clustering Solutions

Selecting the optimal clustering solution remains challenging, as different model-selection criteria often disagree and may point to different solutions, with no principled basis for favoring one over another (Eickhoff et al., 2015; Liu et al., 2020). We therefore based our granularity selection on a heuristic combination of the four following criteria, prioritizing the solution with the greatest convergence across them:

**1. Internal validity metrics:** These assess how similar each subject’s connectivity values correspond to their assigned cluster (i.e., intra-cluster distance) relative to their distinctiveness from other clusters (i.e., inter-cluster distance). The primary measure was the Silhouette index, complemented by the Davies-Bouldin and Calinski-Harabasz indices (see **Supplementary Methods 1** for a detailed description of each measure).
**2. External similarity metrics:** These evaluate the agreement between individual-level cluster labels and the group-level parcellation at each granularity level. We used the adjusted rand index (ARI), adjusted mutual information (AMI) and joint-entropy-normalized variation of information (NVI). AMI and NVI were chosen because their chance- and entropy-based normalization makes them robust to the number of clusters, neither rewarding nor penalizing higher granularity, and thus suitable for comparison across *k* (Kraskov et al., 2005; Vinh et al., 2010). Higher ARI and AMI values and lower NVI values indicate greater alignment between individual- and group-level clustering solutions. These metrics were complemented by relabeling accuracy (see **Supplementary Methods 1** for a detailed description of each measure). Together, these internal and external validity metrics are commonly used to evaluate clustering quality (Eickhoff et al., 2015; Genon et al., 2017; Liu et al., 2021; Reuter et al., 2020).

We analyzed each metric separately using one-way repeated-measures analysis of variance (ANOVA), with granularity as a within-subject factor. If ANOVA revealed a significant main effect of granularity (*p* < 0.05), sequential granularity pairs were compared using paired Student’s *t*-tests, with Bonferroni correction for multiple comparisons.

**3. Cytoarchitectonic similarity:** We assessed the similarity between our TMS-PFC parcellation results and the cytoarchitectonic architecture of DLPFC based on the Jülich-Brain Cytoarchitectonic Atlas (v3.1), which currently includes nine areas covering the middle and superior frontal gyri (Amunts et al., 2020, 2023; Bruno et al., 2024). In addition, we considered the four BA8 subregions, the two Broca’s area subregions, and the yet-to-be-released GapMap Frontal-I because our ROI reached part of these regions. For this, the atlas was converted into MNI152NLin6Asym space with 2 mm isotropic voxels. Only overlapping voxels between the TMS-PFC seed and the cytoarchitectonic atlas were kept (3233 voxels). Next, the ARI was computed between each parcellation result and the overlapping cytoarchitectonic regions. Additionally, we computed the overlap ratio between functional and cytoarchitectonic clusters by dividing the number of overlapped voxels by the total number of voxels in the functional cluster.
**4. Granularities without spatially discontinuous clusters:** To enhance the interpretability of functional connectivity, we prioritized parcellations with spatially continuous clusters (see **Supplementary Methods 2** for the specific criterion to define spatial discontinuity). Ensuring spatial continuity is particularly crucial for potential targeted treatment interventions such as TMS, where fragmented or spatially discontinuous clusters are not only neurobiologically debatable (Bijsterbosch et al., 2020; Eickhoff et al., 2015) but also impractical due to constraints on stimulation precision.

### 2.6 Characterization of Derived TMS-PFC Clusters

Following the selection of our optimal parcellation solution of TMS-PFC, we characterized the resulting TMS-PFC subregions along three complementary levels: SGC connectivity, whole-brain resting-state connectivity profiles, and associated behavioral functions. Since SGC connectivity is the primary anchor of current connectivity-based TMS targeting approaches, we examined it separately across all granularities with spatially continuous clusters, situating our solution within the broader parcellation hierarchy and identifying the consistency of SGC-anticorrelated subregions across granularities. Our primary interest, however, lies in characterizing which large-scale functional networks are reached by clinically used stimulation sites beyond SGC. The whole-brain characterization therefore adopts a multivariate approach, considering each subregion’s overall connectivity pattern across all participants, rather than their connectivity with a single target region.

#### Functional Connectivity with SGC

The SGC was defined based on the Jülich-Brain Cytoarchitectonic Atlas (Palomero-Gallagher et al., 2015), delineating an SGC-ROI by combining areas 25, s24, s32, and the ventral portion of area 33. To isolate the ventral portion of area 33, we limited the ROI to voxels located at or below z = 0, excluding all more dorsal voxels. The resulting SGC-ROI was divided into the left and right hemispheric portions. Functional connectivity values between each TMS-PFC cluster and the left and right SGC were computed for each participant and each level of granularity. To this end, we used voxel-level RSFC matrices from both rs-fMRI sessions, obtained via the rCBP approach, as described in the *Regional Connectivity-Based Parcellation* section. For each participant, we averaged the RSFC values between voxels within the respective TMS-PFC cluster and those within the SGC-ROI.

#### Whole-Brain Functional Connectivity Profiles

Following the selection of our optimal parcellation solution of TMS-PFC, we examined which brain regions were functionally connected with the TMS-PFC subregions to characterize their RSFC profile. We assessed both the overall connectivity pattern of each cluster (main effect) and cluster-specific RSFC profiles using conjunction analysis of pairwise cluster comparisons.

Resting-state fMRI images were preprocessed, as described in the *MRI Preprocessing* section. For each session, individual RSFC estimates were computed by calculating Pearson’s correlation coefficients between the average time series of voxels within each TMS-PFC cluster and the time series of all remaining GM target voxels (156764 voxels), followed by Fisher’s *z*-transformation. The resulting RSFC matrices were averaged across both sessions. We used nilearn (v0.10.4) to enter the *z*-scored RSFC matrices into a group-level mass-univariate generalized linear model (GLM) to compute the main effect of each cluster as well as pairwise cluster comparisons. All contrasts were restricted to the GM mask and thresholded at *p* < 0.05, applying Bonferroni correction for multiple comparisons across all voxels. To derive cluster-specific RSFC profiles, we applied minimum conjunctions across all relevant pairwise cluster comparisons, as well as the main effect of positive or negative RSFC of the respective cluster. For example, for *k* = 3, specific (stronger) positive connectivity of cluster one would be derived by the conjunction of the contrasts C1 > C2, C1 > C3, and the main effect of positive connectivity of C1. Conversely, specific negative connectivity of cluster one would result from the conjunction across the contrasts C1 < C2, C1 < C3, and the main effect of negative connectivity of C1.

To assess the spatial correspondence between cluster RSFC profiles and established large-scale neurocognitive networks, we compared them to 7- and 17-network Yeo atlases (Yeo et al., 2011) using the Network Correspondence Toolbox (Kong et al., 2025). To quantify the magnitude and statistical significance of the alignment between the connectivity maps and network labels, the toolbox first projects the connectivity maps from a standard space into fsaverage6 space, after which correspondence is quantified using the Dice coefficient, where 0 indicates no spatial overlap and 1 indicates complete overlap. The Dice coefficient is calculated as twice the size of the intersection of the two sets divided by the sum of their individual sizes. Statistical significance is then assessed via a spin-test permutation with 1000 rotations for each network (Alexander-Bloch et al., 2018; Vos de Wael et al., 2020). In this procedure, the spherical projection of the cortical surface is rotated randomly 1000 times to generate null models of overlap, with the same rotation applied to every vertex to preserve the spatial features of the original map in a rotated frame. A Dice coefficient is computed for each permutation, and the initially observed Dice coefficient is then compared against the resulting null distribution to determine whether it exceeds the significance threshold of *p* < 0.05.

#### Behavioral Functional Profiling

To infer the behavioral functions associated with the TMS-PFC clusters, we used *behavioral domains* (BDs) and *paradigm classes* (PCs) from the BrainMap database (http://www.brainmap.org) (P. T. Fox & Lancaster, 2002; Genon et al., 2018; Laird et al., 2005, 2011). BrainMap is a curated repository of over 4000 manually encoded task-activation studies that include the peak activation coordinates of brain regions engaged in specific behavioral conditions. These conditions are categorized using an expert-defined ontology, classifying the conditions into BDs, which encompass broad functional categories (e.g., cognition, action, perception, emotion, and interoception) and their related subcategories, and PCs, which denote specific experimental tasks.

We processed each derived cluster of our optimal TMS-PFC clustering solution as follows: First, all experiments in the BrainMap database reporting at least one activation focus within the respective cluster were extracted. Quantitative functional profiling of each TMS-PFC cluster was then achieved via Bayesian *forward inference* and *reverse inference* analyses. *Forward inference* quantifies the probability of observing activity in a given brain region given the presence of a behavioral condition (*P*(Activation|Task)). Conversely, *reverse inference* estimates the probability of a behavioral condition being present, given activation in a specific brain region (*P*(Task|Activation)). For both types of inference, statistical significance was assessed using a binomial test, comparing the conditional probability associated with each BD or PC against its *a priori* base rate across the entire BrainMap database. The base rate is the overall probability of finding activation within the respective cluster (*forward inference*) or the overall prevalence of the given BD or PC (*reverse inference)* in the database. We report both uncorrected results (*p* < 0.05) and results corrected for multiple comparisons using false discovery rate (FDR, *p* < 0.05).

## 3 Results

We first describe the clustering solutions and quantitatively evaluate them to select an optimal granularity based on internal and external validity, cytoarchitectonic overlap, and spatial coherence. The optimal clustering solution was then characterized in terms of its SGC connectivity, whole-brain RSFC profiles, and behavioral associations. We conclude with an overlap map integrating the connectivity between TMS-PFC and SGC across granularities.

### 3.1 Regional Connectivity-Based Parcellation

The parcellation of TMS-PFC based on whole-brain RSFC revealed an anterior-to-posterior division already evident in the two-cluster solution (**Figure 1A**). As granularity increased, this anterior-to-posterior pattern got more pronounced, and a ventral-to-dorsal axis emerged within both anterior and posterior subdivisions. For example, in the three-cluster solution, the anterior cluster two (*k* = 2) was further divided into ventral and dorsal clusters two and three (**Figure 1A**). **Figure 2A** illustrates the hierarchical separation of the clusters across the different granularities. This pattern continued in the four- and five-cluster solutions, in which the anterior section was subdivided into anterior-dorsal, anterior-central, and anterior-ventral clusters two to four, respectively (**Figures 1A and 2A**). Additionally, at *k* = 5, the anterior and posterior sections were further separated by an additional thin cluster two in between them, which, however, was not evident anymore in any other solution. At *k* = 6, the anterior area displayed four clusters along the ventral-to-dorsal axis, whereas the posterior region was further divided into dorsal and ventral clusters one and four. In the seven-cluster solution, the posterior part was split into a single parcel (cluster one) consisting of dorsal and ventral parts as well as an additional ventral parcel (cluster five) lying between the anterior and posterior sections. Additionally, the anterior part comprised five clusters along the ventral-to-dorsal axis, which was also evident in the following granularity (*k* = 8). Moreover, the eight-cluster solution revealed again a division of the posterior region into dorsal and ventral clusters one and four. At *k* = 9, the anterior section retained five clusters along the ventral-to-dorsal axis, whereas the posterior section contained a dorsal cluster one and two ventral clusters five and seven (**Figures 1A and 2A**). Additionally, a small central cluster four emerged between the anterior and posterior regions. Finally, the 10-cluster solution contained six clusters in the anterior region, three clusters in the posterior region, and two small clusters between these two regions.

**Figure 2.**
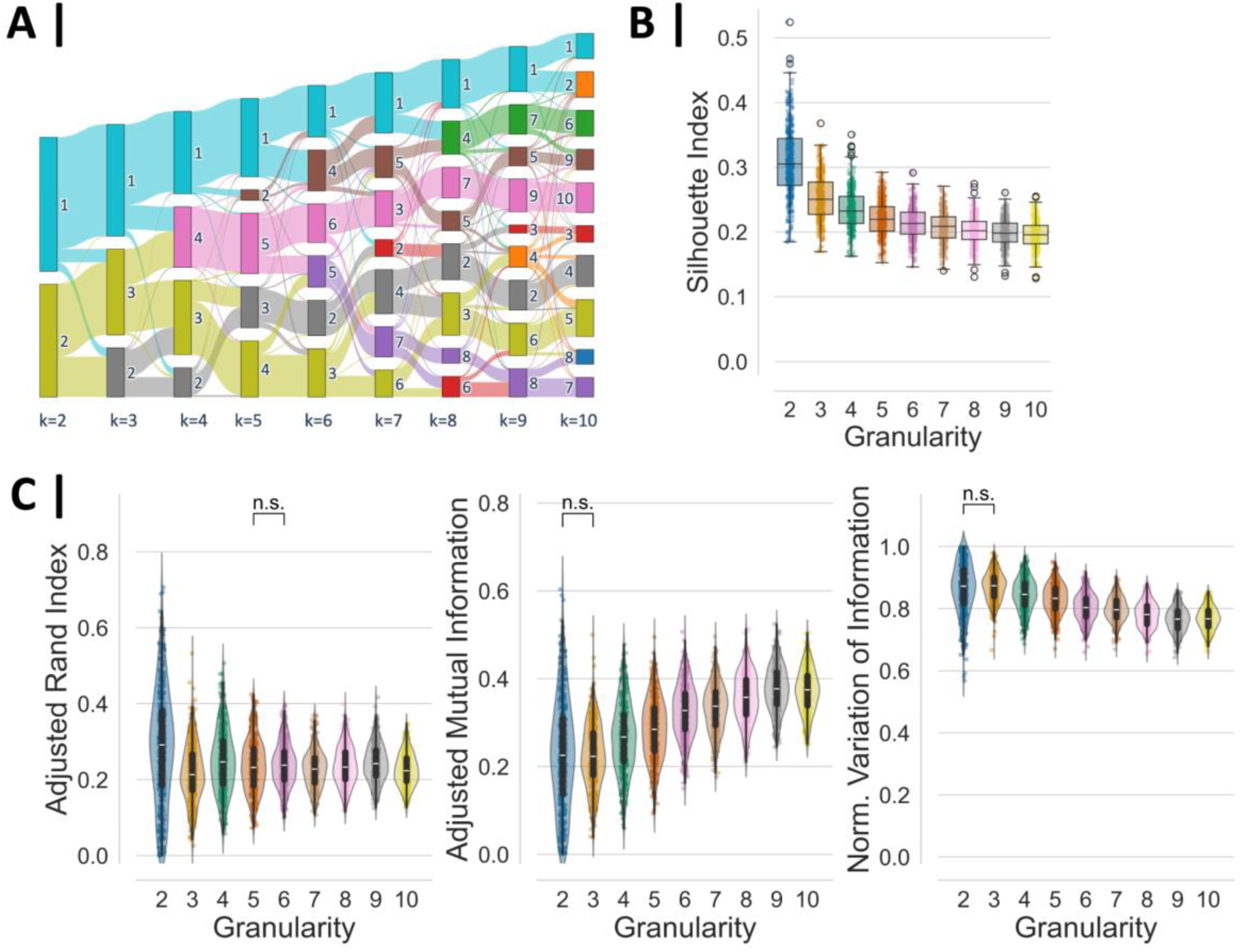
Hierarchical structure and evaluation of the different clustering solutions. **A.** Hierarchical structure of the clustering emerging across granularities. **B.** Internal validity as measured with the Silhouette index. All post-hoc comparisons between sequential pairs with increasing granularity became significant (*p* < 0.001). **C.** External similarity to the reference image for each granularity level, as measured with ARI (left), AMI (center), and NVI (right). All post-hoc comparisons became significant (*p* < 0.001), except for one single pair in each metric marked with n.s. (*p* > 0.05). **Abbreviations.** AMI: adjusted mutual information, ARI: adjusted rand index, n.s.: non-significant, NVI: joint-entropy-normalized variation of information, RSFC: resting-state functional connectivity.

To illustrate the consistency of voxel-wise cluster assignment across participants for each granularity level, we show the corresponding probabilistic maps in **Figure 1B**. Note that to improve interpretability and reduce ambiguity at cluster boundaries, the probabilistic maps were thresholded at 30%, retaining only voxels with sufficient between-subject consistency. Voxels with probabilities below this threshold or those showing spatial overlap between clusters were excluded and labeled as 0, ensuring clearer delineation and facilitating visualization.

At the lowest granularity (*k* = 2), the anterior and posterior emerged as highly consistent clusters, with maximum probability values of 0.95 and 0.96, respectively, indicating strong between-subject agreement in the anterior–posterior division. As the number of clusters increased, the probability of voxel assignment remained relatively high within the core regions of each cluster, particularly in the posterior and anterior-ventral portions of the TMS-PFC, but declined slightly overall. In contrast, peripheral and boundary regions between clusters showed lower probability values. For instance, at *k* = 3, maximum cluster probabilities ranged from 0.63 to 0.81. At *k* = 4, they varied more broadly, from 0.50 to 0.87, indicating increased inter-individual variability at finer parcellation levels. This pattern continued across higher granularities, with consistent but slightly reduced peak probabilities in core subregions (e.g., 0.83 for anterior-central cluster four at *k* = 5, 0.75 for anterior-ventral cluster six at *k* = 6, and 0.71 for anterior-central cluster seven at *k* = 7) and greater heterogeneity along cluster boundaries. From *k* = 8 onward, the highest cluster probabilities remained in a similar range (0.73 to 0.75 for the anterior-ventral clusters at *k* = 8–10), but many peak probabilities dropped below 0.60, suggesting increased inter-individual variability and less stable between-subject alignment. At *k* = 10, the probabilistic maps showed reduced spatial coherence, particularly for some of the smaller subdivisions.

Overall, the probabilistic maps reflect a trade-off between granularity and consistency: Low-*k* solutions yielded broader, more stable clusters with higher between-subject agreement, whereas higher-*k* solutions offered more fine-grained parcellations with slightly reduced spatial coherence across participants. This pattern reflects both increased inter-individual variability and the inherent association between lower probabilities and higher *k-*values. Nevertheless, several clusters still exhibited peak probabilities around 0.7, reflecting a reasonably robust level of consistency even at higher granularity levels.

In sum, the TMS-PFC is organized along two principal axes: A stable anterior-posterior division and a ventral-dorsal gradient within each subdivision. This hierarchical structure provides a balance between broad organizational stability and finer-grained differentiation that emerges at higher parcellation levels.

### 3.2 Evaluation of Clustering Solutions

Following the approach described in the methods, we systematically evaluated each cluster solution according to the internal and external validity, correspondence with a cytoarchitectonic map, and spatial coherence, prioritizing the solution with the greatest convergence across all criteria.

#### Internal and External Validity

The Silhouette index (**Figure 2B**), as well as the Davies-Bouldin and Calinski-Harabasz indices (**Supplementary Results, Figure S1A**), displayed a nonlinear monotonic decline (increase for the Davies-Bouldin Index) in their mean and variance values as the number of clusters increased, with a Silhouette index of 0.31 ± 0.05 at *k* = 2 and reaching the lowest value at *k* = 10 with 0.20 ± 0.02. The substantial effect of granularity was confirmed by a one-way repeated-measures ANOVA, *F*(8,3120) = 1325.41, *p* < 0.001, *η_p_^2^* = 0.25. All post-hoc comparisons of sequential pairs with increasing granularity showed a significant decline (*p* < 0.001) after multiple-comparisons correction.

It is worth noting that low-granularity solutions such as *k* = 2 are inherently favored by internal validity metrics, as fewer, larger clusters produce greater intra-cluster homogeneity and inter-cluster separation by construction (Liu et al., 2020; Plachti et al., 2019; Reuter et al., 2020). This represents a well-known limitation of these metrics rather than evidence of functional neurobiological optimality, underscoring the need for complementary criteria to identify a more informative granularity (Eickhoff et al., 2015).

The ARI revealed the highest similarity between individual-level and group-level clusterings for *k* = 2 (ARI*_k=2_* = 0.29 ± 0.14; **Figure 2C**), followed by the four- and nine-cluster solutions (ARI*_k=4_* = 0.25 ± 0.08 and ARI*_k=9_*= 0.24 ± 0.05). The ANOVA confirmed a significant main effect of granularity, *F*(8,3120) = 53.06, *p* < 0.001, *η_p_^2^* = 0.28. Nevertheless, the trajectory of the ARI values was non-monotonic, showing an initial sharp decline from *k* = 2 to 3, followed by an increase at *k* = 4. Subsequently, ARI values continued to decrease until *k* = 7, at which point they significantly increased for *k* = 8 and 9 before dropping again in the 10-cluster solution. These fluctuations suggest that certain granularities (*k* = 2, 4, 8, 9) preserve individual-to-group similarity better than others, with *k* = 9 representing a local maximum after *k* = 2 and *k* = 4. It should be noted, however, that the higher absolute values at *k* = 2 and *k* = 4 were accompanied by properties that limit their suitability as a final parcellation: The two-cluster solution showed the largest inter-subject variability (SD = 0.14 versus 0.05 at *k* = 9), and the four-cluster solution contained spatially discontinuous parcels (see *Spatial Coherence* below).

In contrast to the ARI, the AMI and NVI showed a broadly monotonic improvement with increasing granularity, consistent with their chance- and entropy-based normalization being robust to the number of clusters. AMI increased from AMI*_k=2_* = 0.23 ± 0.12 to the highest value at *k* = 9 (AMI*_k=9_* = 0.38 ± 0.05; **Figure 2C**), revealing higher information shared between individual- and group-level clusterings with increasing granularity. Supporting this finding, NVI decreased from NVI*_k=2_* = 0.87 ± 0.08 to the lowest values at *k* = 9 (NVI*_k=9_* = 0.77 ± 0.04), revealing a reduction in leftover uncertainty about one clustering given knowledge of the other. Both metrics confirmed a significant main effect of granularity (AMI: *F*(8,3120) = 678.10, *p* < 0.001, *η_P_^2^* = 0.31; NVI: *F*(8,3120) = 721.10, *p* < 0.001, *η_P_^2^* = 0.33). Post-hoc comparisons of sequential granularity pairs were significant after multiple-comparison correction, except for *k* = 2 versus *k* = 3 for both metrics. Notably, this trend reversed at the highest granularity tested: Both AMI and NVI deteriorated significantly from *k* = 9 to *k* = 10 (AMI: *t*(390) = 3.69, *p* = 0.009; NVI: *t*(390) = −3.39, *p* = 0.028), identifying *k* = 9 as the optimum within the tested range. Notably, the between-subject variability was largest for the two-cluster solution and progressively decreased as granularity increased, indicating more consistent individual-to-group correspondence at finer granularities. The results for relabeling accuracy are provided in the **Supplementary Results (Figure S1B)**.

While the internal metrics point to low-*k* solutions, both AMI and NVI identified *k* = 9 as the granularity with the highest individual-to-group similarity, with a small but significant deterioration at *k* = 10. Together with the local ARI maximum at the same granularity, this consistency across three similarity measures supports *k* = 9 as the most robust clustering solution among those tested.

#### Cytoarchitectonic Similarity

To examine the correspondence of our TMS-PFC parcellation with the underlying cytoarchitectonic organization, we compared it with the nine DLPFC areas defined in the Jülich-Brain Cytoarchitectonic Atlas (v3.1) and surrounding overlapping areas (Amunts et al., 2020, 2023; Bruno et al., 2024), acknowledging that connectivity-based parcellations do not necessarily follow cytoarchitectonic boundaries. The comparison showed that the correspondence strongly increased from *k* = 3 to 4 and reached the highest values with some variability between *k* = 8, 9, and 10 (see **Supplementary Results, Figure S2 and Table S2**).

#### Spatial Coherence

To assess the spatial coherence of the clustering solutions, we examined whether clusters contained spatially discontinuous components larger than five voxels under the same label. Only the two-, three-, six-, eight-, and nine-cluster solutions met this criterion, indicating consistent and continuous spatial organization. In contrast, other granularities exhibited discontinuities, with 25 voxels of spatially discontinuous components in the four-cluster solution, 43 voxels at *k* = 5, 12 voxels in the seven-cluster solution, and 180 voxels at *k* = 10.

Taken together, these analyses converge on *k* = 9 as the optimal granularity. While low-*k* solutions provided stable and broad divisions and are favored by the internal validity metrics, they showed comparatively low cytoarchitectonic overlap and lower correspondence between individual- and group-level clustering. Among the higher granularities, *k* = 10 exhibited the largest spatial discontinuity (180 voxels for one cluster), and *k* = 8, despite adequate spatial coherence and cytoarchitectonic correspondence, did not reach the individual-to-group similarity observed at *k* = 9. The nine-cluster solution was the only granularity that combined the highest external validity (ARI local maximum, highest AMI, lowest NVI), among high cytoarchitectonic correspondence, and full spatial coherence. We therefore selected this level of granularity as the optimal parcellation solution to represent the TMS-PFC and for the subsequent characterization analyses. For the separate probabilistic maps of each cluster within the nine-cluster solution, please refer to the **Supplementary Results, Figure S3.**

### 3.3 Characterization of TMS-PFC Nine-Cluster Solution

We next characterized the nine-cluster solution in terms of its SGC connectivity, its large-scale connectivity patterns, and its behavioral associations to gain a comprehensive understanding of the TMS-PFC’s functional architecture.

#### Functional Connectivity with SGC

SGC connectivity is the main anchor of current connectivity-based TMS targeting approaches in TRD. We therefore examined it separately across granularities with spatially continuous clusters (*k* = 2, 3, 6, 8, 9), assessing the consistency of SGC-connected subregions within the broader parcellation hierarchy.

In the two-cluster solution, the anterior cluster (*k*-clusterID: 2-2) exhibited negative functional connectivity with the bilateral SGC (Fisher’s *z* = −0.11 ± 0.07), whereas the posterior cluster (2-1) showed positive connectivity (Fisher’s *z* = 0.10 ± 0.07; **Figure 3**). As granularity increased (*k* = 3, 6, 8), this pattern became more refined, with an anterior-central cluster (3-3, 6-3, and 8-3, see **Figure 3**) consistently remaining the most anti-correlated with SGC, while the posterior (3-1) and superior-posterior clusters (6-1 and 8-1) showed the opposite pattern. At *k* = 9, the two extremes were most clearly delineated: The anterior-central cluster (9-6) exhibited the strongest functional anti-correlation with SGC (Fisher’s *z* = −0.13 ± 0.07), while the superior-posterior cluster (9-1) displayed the strongest positive connectivity (Fisher’s *z* = 0.16 ± 0.08).

**Figure 3.**
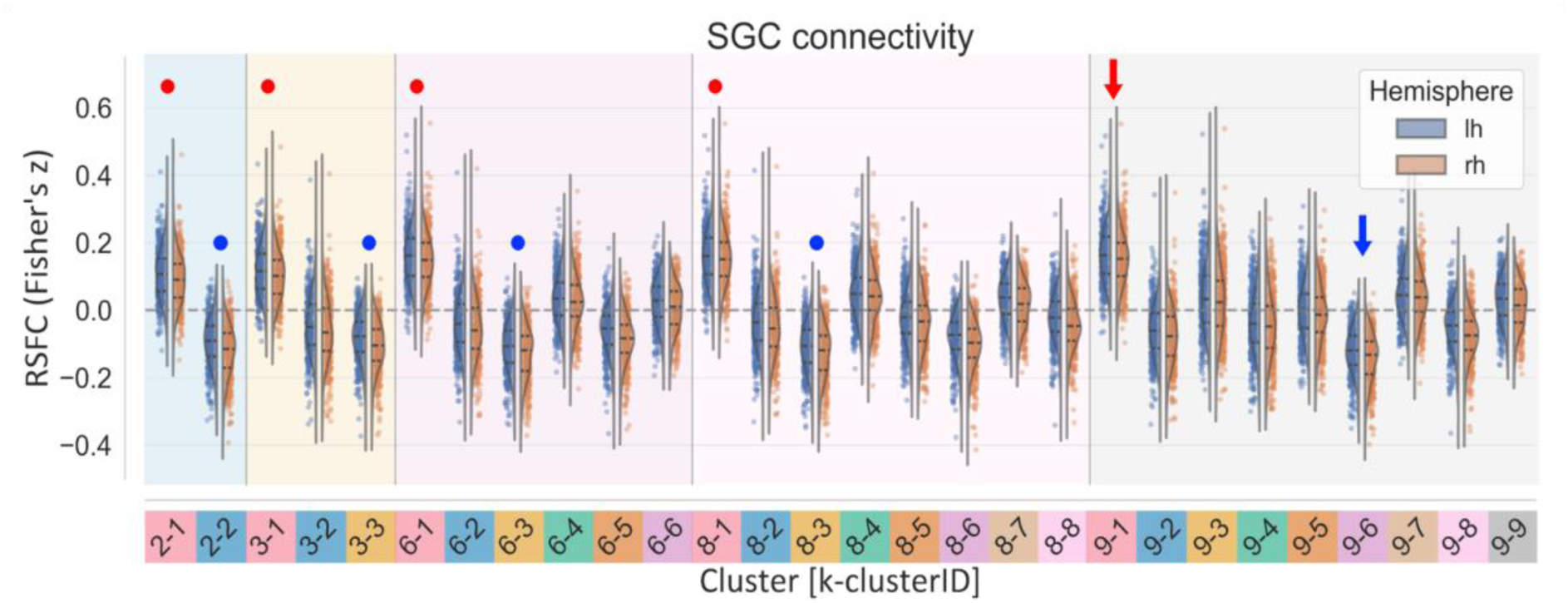
SGC functional connectivity for the TMS-targeted left prefrontal cortex subregions across granularities. RSFC between each cluster and left and right SGC for granularities with spatially continuous regions. Red and blue dots mark the cluster within each granularity with the strongest positive and negative functional connectivity with SGC, respectively. The arrows mark the superior-posterior (cluster 1) and anterior-central (cluster 6) clusters at *k* = 9 that maximized the positive and negative RSFC with bilateral SGC, respectively. **Abbreviations.** RSFC: resting-state functional connectivity, SGC: subgenual cingulate cortex, TMS: transcranial magnetic stimulation.

This anterior-to-posterior antagonism was thus preserved across all spatially coherent granularities, with the anterior-central subregion consistently being the most SGC-anticorrelated portion of TMS-PFC and the posterior-superior subregion consistently being the cluster with the positive SGC connectivity. Notably, finer granularities also isolated subregions with negligible SGC connectivity (e.g., clusters 9-3, 9-5, and 9-9; **Figure 3**), which at coarse granularities were absorbed into larger clusters showing non-zero mean connectivity. Thus, increasing granularity not only refined the anterior-to-posterior organization but also delineated subregions that carry little SGC-related information.

Beyond the single ROI-based SGC connectivity analysis, we next examined the whole-brain connectivity profile of each TMS-PFC subregion in the nine-cluster solution.

#### Resting-State Functional Connectivity Profiles

When assessing the RSFC profiles of the nine-cluster solution, we observed that posterior TMS-PFC subregions (clusters one, five, and seven) exhibited a broadly similar connectivity pattern (**Figure 4A**). All of them showed strong positive connectivity with regions overlapping with the default mode network (DMN) as well as functional anti-correlation with the salience network (SAN). Posterior-ventral parts (clusters five and seven) additionally showed negative functional connectivity with the somatomotor (SMN) and visual networks (VIS), and posterior clusters one and seven with the dorsal attention network (DAN). Central-ventral cluster five’s positive connectivity map also overlapped with the frontoparietal network (FPN).

**Figure 4.**
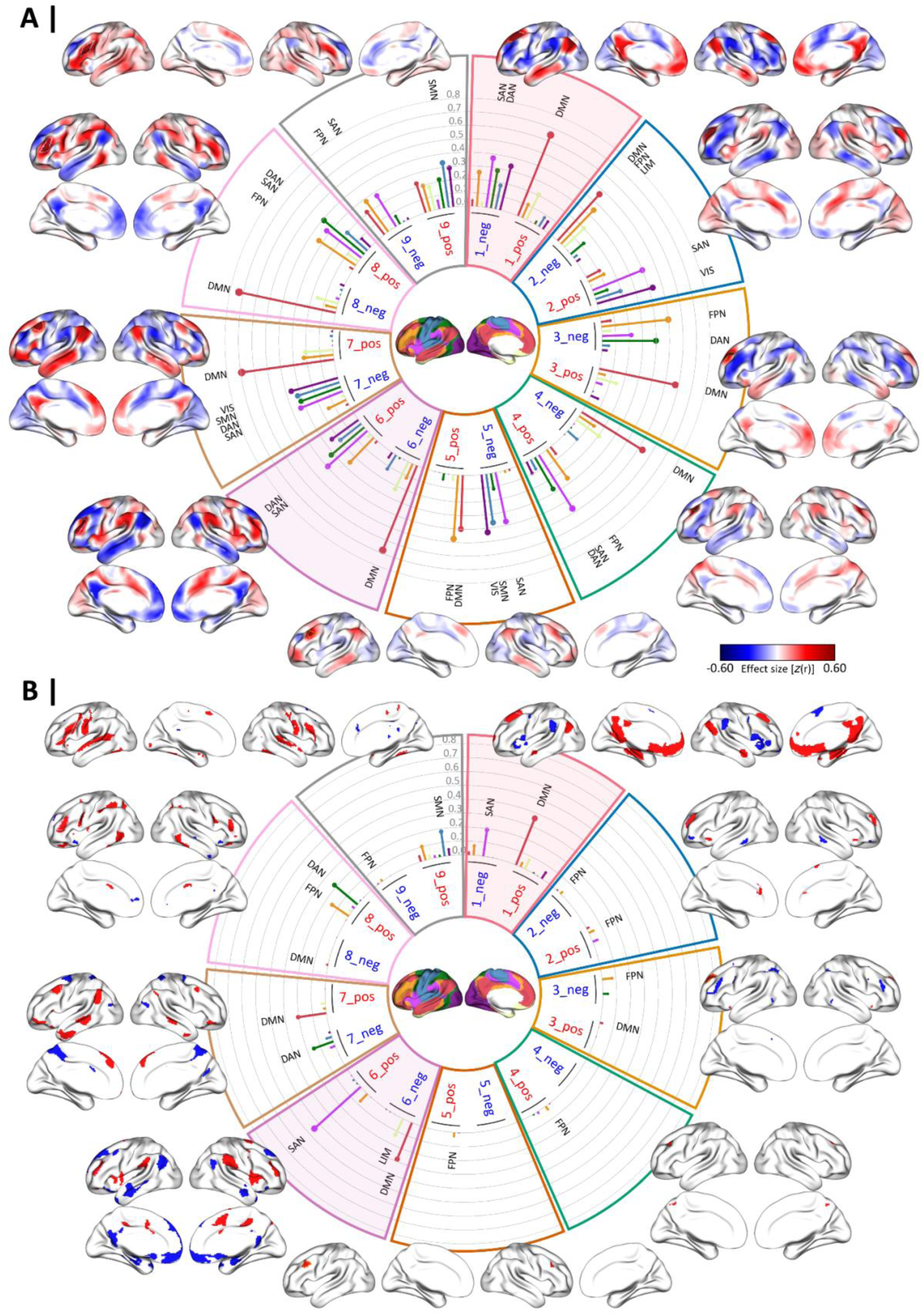
Resting-state functional connectivity profiles of the TMS-targeted left prefrontal cortex subregions. **A.** Positive and negative RSFC main effects for each cluster of the nine-cluster solution. **B.** Cluster-specific RSFC assessed via conjunction of pairwise comparisons for each cluster of the nine-cluster solutions. The circular charts illustrate the correspondence (Dice coefficients) between the 7-Yeo networks and the positive and negative RSFC maps, separately. Labeled bars denote statistically significant overlap larger than expected by chance after spin-test permutations. The shaded sectors emphasize the clusters with strongest positive (cluster one) and negative (cluster six) connectivity with the subgenual cingulate cortex. **Abbreviations.** DAN: dorsal attention network, DMN: default-mode network, FPN: frontoparietal network, LIM: limbic network, neg: negative, pos: positive, RSFC: resting-state functional connectivity, SAN: salience attention network, SMN: somatomotor network, TMS: transcranial magnetic stimulation, VIS: visual network.

In contrast, the anterior-central clusters six and eight, as well as the centrally located cluster four, directly posterior to them, exhibited roughly the opposite pattern: For instance, clusters four, six, and eight showed functional anti-correlation with DMN, while displaying positive connectivity with SAN and DAN (**Figure 4A**). The positive RSFC of clusters four and eight further overlapped with FPN. Cluster two, situated dorsally to cluster six, was also positively connected with SAN and VIS, negatively associated with DMN and the limbic network (LIM), but also negatively connected with regions of FPN. The small anterior-dorsal cluster three displayed a pattern mainly akin to the posterior subregions, characterized by positive connectivity with DMN and anti-correlations with FPN and DAN. The ventral cluster nine presented a distinct profile, marked by positive RSFC with SMN and anti-correlations with SAN and FPN.

Beyond this principal antagonism, clusters seven, eight, and nine also exhibited notable cluster-specific connectivity profiles. While cluster eight shared the broadly anterior-central pattern of cluster six, its cluster-specific profile was distinguished by positive connectivity primarily with DAN and FPN regions, particularly in the inferior posterior temporal gyrus, superior parietal lobule (SPL), and ventral prefrontal regions, distinguishing it from cluster six (**Figure 4B**). Cluster seven, despite broadly following the posterior cluster one pattern of positive DMN connectivity, showed more distinguished cluster-specific negative connectivity with DAN, mainly with the precuneus, suggesting a partially distinct posterior specialization. Cluster nine showed cluster-specific connectivity concentrated around primary somatomotor regions and superior temporal gyrus. While clusters two, three, four, and five showed more constrained cluster-specific connectivity profiles, they still exhibited slightly different profiles based on RSFC of their main effects (**Figure 4A**), suggesting they differ in their functional profile but with no region that is consistently different compared to all other subregions. The network correspondence for these RSFC main effects and cluster-specific RSFC maps with the 17-Yeo networks is provided in the **Supplementary Results (Figure S4**).

In summary, our findings revealed a two-partition within TMS-PFC in terms of an antagonistic pattern of RSFC with DMN vs. SAN and DAN: Posterior regions (clusters one, seven, and five) being positively connected with DMN and negatively with SAN and DAN, whereas anterior-central subregions (clusters six, eight, and four) exhibited the opposite pattern. Importantly, while multiple clusters contributed to this broad antagonism, clusters one and six maximized the specific connectivity to these networks when compared to all other clusters (**Figure 4B**), representing the most differentiated poles in this organization. The finer cluster-specific differences, particularly between clusters one and seven posteriorly, and clusters six and eight anteriorly, further underscore the functional heterogeneity within TMS-PFC, expanding our understanding of its organization.

#### Behavioral Functional Profiling

When assessing the behavioral domains associated with TMS-PFC subregions from our selected cluster solution (*k* = 9) using BrainMap, we observed a mixed pattern of domains associated with each cluster. Anterior clusters, including clusters six and eight, but also (posterior) cluster five, were significantly associated with working memory, whereas more posterior regions, such as clusters one and seven, were primarily engaged in explicit memory (**Figure 5**). Along the ventral-to-dorsal axis, the ventral-most cluster (cluster nine) was primarily linked to language-related functions, consistent with its proximity to Broca’s areas, and also showed associations with both working and explicit memory. In contrast, dorsal regions, including clusters two and three, were significantly involved in social cognition. Notably, posterior cluster seven also showed involvement in social cognition. Central cluster four showed significant associations with action inhibition, as did the clusters anterior to it, clusters six and two, although these did not reach significance. Additionally, centrally located cluster four (and cluster five, although not significantly) was involved in color perception. Interestingly, cluster two was the only cluster significantly associated with thermoregulation and pain perception.

**Figure 5.**
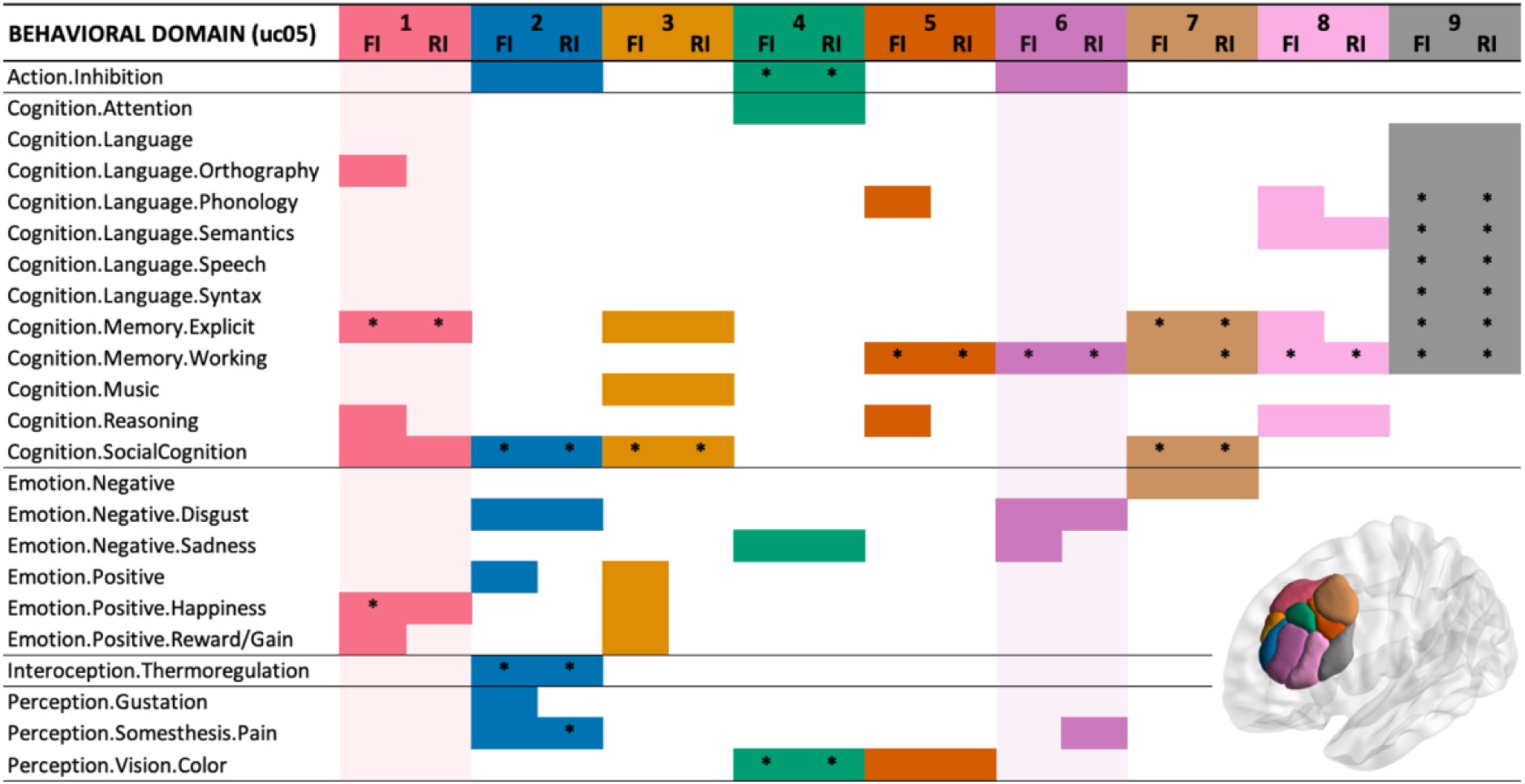
Behavioral functional profiling of the TMS-targeted left prefrontal cortex subregions. Behavioral domains associated with each cluster of the nine-cluster solution. The shaded sectors emphasize the clusters with strongest positive (cluster one) and negative (cluster six) connectivity with the subgenual cingulate cortex. No significant associations behind the brain image. **Abbreviations.** FI: forward inference as P(Activation|Task), RI: reverse inference as P(Task|Activation), TMS: transcranial magnetic stimulation, uc05: uncorrected *p* < 0.05, * FDR-corrected *p* < 0.05.

With regard to behavioral domains of emotions, distinct patterns emerged between clusters, although only at the uncorrected level. Of particular relevance for depression, anterior-central cluster six showed engagement in processing negative emotions such as disgust and sadness, which is partly shared with clusters two, four, and seven. In contrast, more dorsal clusters (clusters one to three) were associated with positive emotions, including happiness and reward. Clusters five, eight, and nine showed no associations with emotional behavioral domains. For an overview of the paradigm classes associated with each of the nine TMS-PFC clusters, please refer to the **Supplementary Results (Figure S5).**

These behavioral profiles supported the network-level dissociation and functional divergence, with posterior clusters linked to explicit memory and positive emotions, anterior clusters to working memory and action inhibition, and ventral and dorsal clusters to language and social cognition, respectively.

### 3.4 TMS-PFC Overlap Map Across Granularities According to SGC Connectivity

Having observed that the antagonistic SGC connectivity pattern was preserved across all spatially continuous granularities, we next delineated the area of TMS-PFC that consistently showed this pattern irrespective of the clustering solution. For each granularity without discontinuous parcels (*k* = 2, 3, 6, 8, 9), we identified the clusters most anti-correlated with SGC (*k*-clusterID: 2-2, 3-3, 6-3, 8-3, 9-6; see **Figure 3**), all consistently located in the anterior portion of TMS-PFC. The positively correlated clusters were all located in the posterior TMS-PFC (cluster one across all granularities). For each connectivity direction, we created a TMS-PFC overlap map via a minimal conjunction across the binary cluster maps of the included granularities. That is, a voxel was assigned to the SGC-region only if it belonged to the most anti-correlated cluster at every included granularity, and to the SGC+ region only if it belonged to the most positively correlated cluster at every granularity (**Figure 6A**). This yielded the TMS-PFC portions whose SGC connectivity profile is independent of the chosen granularity. The SGC+ and SGC-areas were then further analyzed in terms of their whole-brain functional connectivity and behavioral associations.

**Figure 6.**
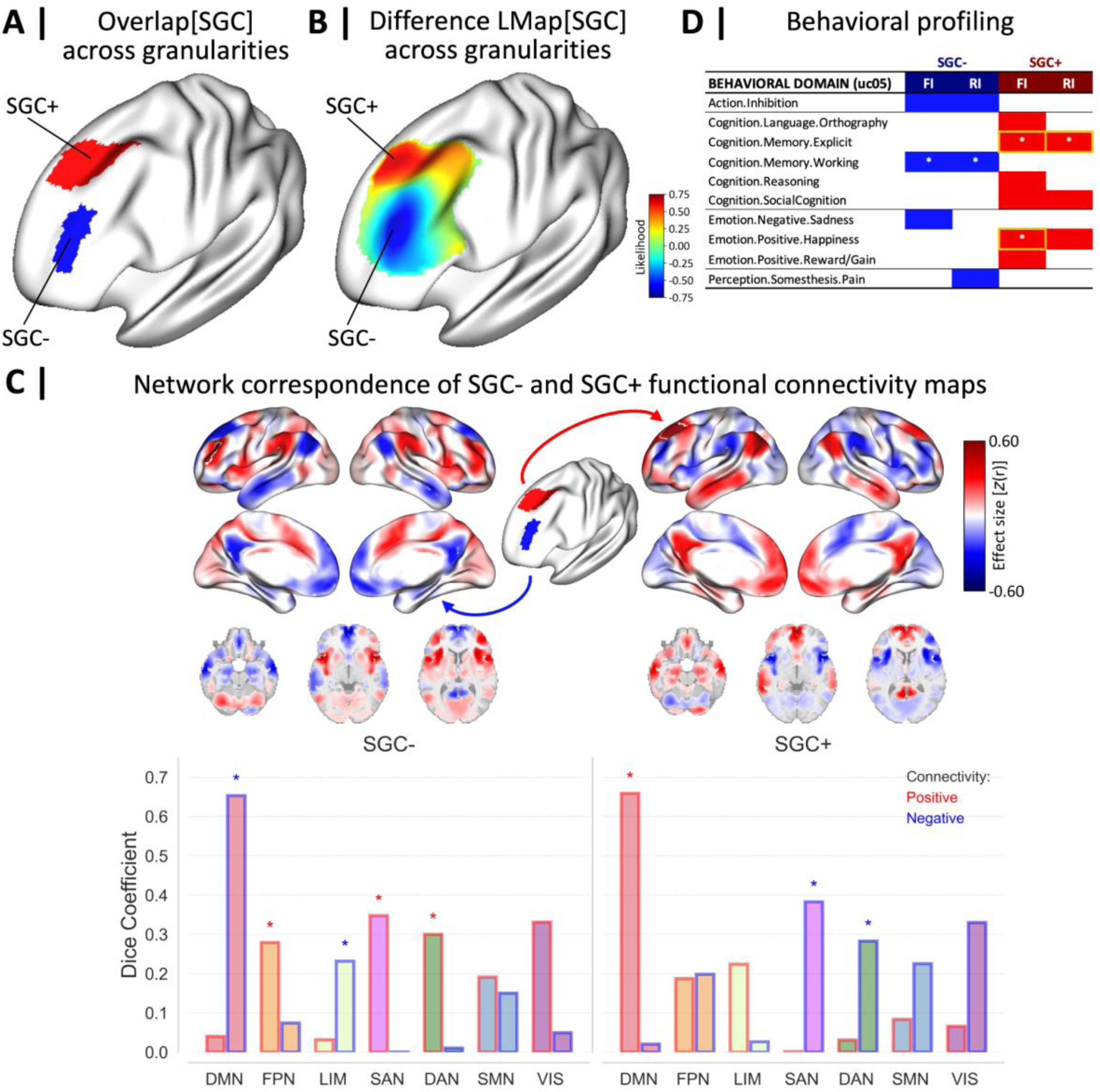
Antagonistic TMS-PFC subregions SGC+ and SGC- and their resting-state functional connectivity and behavioral profiles. **A.** Overlap (minimal conjunction) of the clusters most positively (SGC+) and most negatively (SGC-) connected with SGC across the same granularities (*k* = 2, 3, 6, 8, 9). **B.** For comparison, difference likelihood map of TMS-PFC subregions according to their connectivity with SGC across granularities, obtained by averaging between-subject cluster probability maps (see **Supplementary Methods 2**). **C.** Positive and negative RSFC main effects for SGC- and SGC+ clusters (panel A) and their network correspondence (Dice coefficients) with the 7-Yeo networks. Bars with asterisks denote statistically significant overlap larger than expected by chance after spin-test permutations. **D.** Behavioral domains associated with SGC- and SGC+ clusters (panel A). The orange frames mark the significant domains (FDR-corrected) that also turned significant when analyzing the contrast between the two clusters. **Abbreviations.** DAN: dorsal attention network, DMN: default-mode network, FI: forward inference as P(Activation|Task), FPN: frontoparietal network, LIM: limbic network, neg: negative, pos: positive, RI: reverse inference as P(Task|Activation), RSFC: resting-state functional connectivity, SAN: salience attention network, SGC: subgenual cingulate cortex, TMS: transcranial magnetic stimulation, TMS-PFC: TMS-targeted left prefrontal cortex, uc05: uncorrected *p* < 0.05, * FDR-corrected *p* < 0.05.

For comparison, we additionally computed a probability-weighted version of this analysis by averaging the between-subject cluster probability maps across granularities and subtracting the SGC-from the SGC+ composite, yielding a difference likelihood map (**Figure 6B**; see **Supplementary Methods 2** for details). Unlike the conjunction, this map is continuous and reflects the graded between-subject consistency of cluster assignment, illustrating that the antagonistic organization extends beyond the strictly overlapping areas as a spatial gradient across the ROI. The two approaches converged strongly: When thresholding the likelihood map at ±50%, SGC+ and SGC-regions and their whole-brain connectivity and behavioral profiles were highly similar to those obtained by the minimal conjunction (**Supplementary Results, Figure S6**). Therefore, we here only present the connectivity and behavioral profiles of the conjunction-defined SGC- and SGC+ regions (**Figure 6A**).

The network correspondence analysis using Dice coefficients revealed distinct profiles for SGC- and SGC+. The negative RSFC profile of SGC-showed significant overlap with regions of DMN and LIM, while its positive RSFC pattern overlapped with SAN, DAN, and FPN (**Figure 6C**, left). Conversely, SGC+ displayed the opposite pattern, with positive connectivity to DMN and negative connectivity to SAN and DAN (**Figure 6C**, right). For additional details regarding the overlap with the 17-Yeo networks, see **Supplementary Results (Figure S7A)**.

Behavioral functional profiling using BrainMap further supported this functional dichotomy (**Figure 6D**). SGC+ showed significant associations with explicit memory and positively valenced emotions of happiness, which remained significant in the statistical contrast against SGC-(**Supplementary Results, Figure S7C)**. At the uncorrected level, SGC+ was also related to positively valenced emotions of reward, social cognition, reasoning, and orthographic processing. In contrast, SGC-showed significant associations with working memory, and at the uncorrected level, with action inhibition, sadness, and pain perception. For the corresponding paradigm classes and contrasts, refer to **Supplementary Results (Figures S7B and S7C)**.

Together, these findings highlight a functional dissociation between SGC+ and SGC-: The former was linked to explicit memory and happiness, whereas the latter was associated with working memory and, at a subthreshold level, with cognitive control and negative emotional processes. This antagonistic pattern mirrors the distinct network affiliation of clusters six and one of the nine-cluster solution, further emphasizing their role in shaping the TMS-PFC’s functional organization.

## 4 Discussion

The present study aimed to identify functionally distinct subregions encompassed by clinically used TMS targets within the left prefrontal cortex (TMS-PFC), based on whole-brain RSFC patterns, and to characterize their SGC connectivity, large-scale network embedding, and associated behavioral profiles. Rather than aiming for a general-purpose parcellation of the lDLPFC, our goal was to establish a functionally meaningful subdivision of the area covered by clinically used stimulation sites, determining which functional subregions these targets reach and how they differ in connectivity and behavioral relevance. In doing so, our study provides an atlas of the TMS-PFC functional organization in healthy young adults, from which testable hypotheses can be derived for future clinical and neuromodulation studies in TRD.

Our findings suggest that the TMS-PFC can be understood at two complementary levels of hierarchy. At a coarse level, it is organized into two antagonistic poles: An anterior-central subregion (SGC-) and a superior-posterior subregion (SGC+). At a finer level, however, the TMS-PFC can be subdivided into nine functionally distinct clusters that expand upon this antagonism. Notably, although lower cluster solutions (e.g., *k* = 2) offered high overall validity, they obscured substantial inter-individual heterogeneity. In contrast, the nine-cluster solution captured more nuanced, behaviorally meaningful subdivisions, highlighting the intrinsic heterogeneity of TMS-PFC and its complex embedding within large-scale brain networks (DMN, FPN, SAN, DAN, LIM, and SMN). This hierarchical view integrates coarse organization and subregional precision: The two-pole framework recapitulates the anterior-posterior antagonism underlying SGC-centric targeting approaches, whereas the nine-cluster parcellation refines this organization by delineating specific network- and behaviorally-defined subregions, offering a finer-grained reference from which hypotheses about clinical implications, including symptom-specific or individually optimized targeting, can be formulated and tested.

### 4.1 TMS-PFC Connectivity-Based Parcellation

In terms of anatomical subdivision, the two-cluster solution resembles traditional cytoarchitectonic divisions (Brodmann, 1909; Sarkissov et al., 1955; von Economo & Koskinas, 1925) and bears similarity to prior task-based subdivisions of the right DLPFC (Cieslik et al., 2013), as well as two orthogonal rostrocaudal and dorsoventral gradients recently identified through meta-analytic decoding (Abdallah et al., 2022). However, the nine-cluster solution provided the clearest and most differentiated picture of TMS-PFC functional organization. The spatial distribution of these clusters aligns more closely with more recent evidence supporting finer-grained subdivisions based on cytoarchitecture, as well as structural and functional connectivity (Bruno et al., 2024; Jung et al., 2022; Sallet et al., 2013). For instance, the subdivision by Jung et al. (2022) based on Brodmann-area-based spherical ROIs also follows an anterior-posterior and ventral-dorsal pattern similar to our higher-order cluster solutions.

Crucially, unlike existing parcellations of (DL)PFC that were all not designed with TMS targets in mind and additionally focus on the right hemisphere (Cieslik et al., 2013; Sallet et al., 2013), rely on cytoarchitectonically defined spherical ROIs (Jung et al., 2022; Li et al., 2024), or use small samples (Sallet et al., 2013), our study is anchored to the spatial distribution of actual clinical TMS coordinates and directly characterizes the subregional heterogeneity within a region of TMS stimulation sites. Specifically, our nine-cluster solution revealed systematic differences in connectivity with SGC and depression-relevant networks, alongside behavioral associations in memory, cognitive control, language, and emotional processes. These clusters not only recapitulated an anterior–posterior antagonism but also revealed distinct ventral and dorsal specializations, highlighting that the TMS-PFC supports a broader spectrum of cognitive, affective, and sensorimotor functions than suggested by coarser subdivisions. Together, these findings establish the nine-cluster solution as a fine-grained framework that aligns with prior anatomical models while informing about connectivity patterns that are relevant to current targeting approaches.

#### Functional Connectivity with SGC

Building on this framework, we examined how TMS-PFC subregions relate to SGC, a central target of connectivity-based TMS treatments for TRD. The broad anterior-posterior distinction we observed in SGC connectivity across TMS-PFC subregions is consistent with prior clinical and neuroimaging evidence. Herbsman et al. (2009) demonstrated, through retrospective TMS outcomes, that more anterolateral stimulation coordinates were associated with better antidepressant efficacy, a coarse clinical observation that lacked mechanistic explanation. M. D. Fox et al. (2012) subsequently showed that this advantage could be explained by stronger SGC anti-correlation at anterolateral areas. Notably, our nine-cluster solution refines this picture substantially: Extending previous studies reporting spatial heterogeneity in RSFC between DLPFC and SGC (Cash, Cocchi, Lv, Wu, et al., 2021; Cieslik et al., 2013; M. D. Fox et al., 2012; Siddiqi et al., 2021), we demonstrate that not all anterior TMS-PFC subregions are equivalently anti-correlated with SGC, a distinction not resolved by coarser parcellations. Specifically, only a subset of anterior-central clusters, namely cluster six, exhibited strong anti-correlation with SGC, whereas other anterior subregions showed weak or negligible connectivity. If this organization holds in the clinical population, it would imply that stimulation of different anterior areas might have different effects on SGC-related circuits, a prediction worth testing against coordinates and treatment outcomes in future clinical studies. Relatedly, the spatial scale of these findings implies that millimeter-scale deviations in coil placement could shift stimulation to functionally distinct subregions, which has been proposed as a potential source of variability in clinical response (Ning et al., 2019). Thus, our work refines previous findings by providing a more fine-grained and differentiated picture of SGC connectivity within TMS-PFC across different levels of granularity (**Figure 3** for reference). Consistent with this, Siddiqi et al. (2021) demonstrated that connectivity to a broader causal depression circuit, extending beyond SGC by including distributed cortical and subcortical areas, predicted TMS outcomes more accurately than SGC connectivity alone. This further underscores the necessity of characterizing the full network embedding of TMS-PFC subregions, which is the focus of the following section.

#### Whole-Brain Functional Connectivity and Behavioral Profiles

Beyond SGC connectivity, our study further provides a multivariate whole-brain characterization of each cluster’s RSFC profile. While the systematic differences in SGC connectivity situate our parcellation within the established SGC-centric framework, our primary interest lies in what these subregions reveal beyond this single seed-based relationship. This was motivated by extensive evidence of large-scale circuit-level alterations in depression, particularly involving DMN, SAN, DAN, and fronto-limbic networks (Anderson et al., 2016; Drysdale et al., 2017; Northoff & Hirjak, 2024). Our results revealed that the anterior-central part of TMS-PFC (clusters six and eight, as well as SGC-in the overlap map) showed negative RSFC with DMN and limbic regions and positive connectivity with SAN, DAN, and FPN (**Figures 4B and 6C** for reference). By contrast, most posterior subregions displayed the inverse pattern. This pattern was most prominent for the anterior-central and superior-posterior SGC- and SGC+ in the overlap map (**Figure 6A** for reference). These findings are consistent with prior evidence identifying the dorsal-posterior DLPFC as a functional node of DMN (Alves et al., 2019), as well as reports of anti-correlation between anterior and posterior lDLPFC areas and antagonistic connectivity with respect to the DMN (Reid et al., 2016).

### Anterior-central areas

Importantly, although most clusters showed some degree of connectivity with parts of the DMN, FPN, SAN, and DAN networks, only a subset, namely the anterior-central clusters (four, six, and eight), fully reflected the canonical triple-network profile (DMN, FPN, SAN) proposed as critical for adaptive cognitive, emotional, and behavioral processing (B. Menon, 2019; V. Menon & D’Esposito, 2022). Cluster six in particular showed the strongest specific anti-correlation with DMN and LIM and specific positive connectivity with SAN, especially TPJ, insula, preSMA, and MCC, distinguishing it from surrounding subregions. Notably, while inferiorly located cluster eight shared the broadly anterior-central triple-network pattern, its cluster-specific profile was characterized by positive connectivity with FPN and DAN rather than SAN, reflecting a partially distinct functional specialization that will be discussed further below. The functional heterogeneity of anterior-central profiles is further underscored by the fact that anterior-superior cluster two and anterior-ventral cluster nine showed no cluster-specific overlap with SAN or DMN, and the superior cluster three exhibited anticorrelation with FPN (**Figure 4** for reference).

These profiles may reflect a role of cluster six in mediating interactions between SAN, as a network that detects and prioritizes salient internal or external stimuli, and DMN, which supports self-referential and social cognitive processes (Seeley et al., 2007; Sridharan et al., 2008; Uddin, 2015). Ample evidence suggests that SAN mediates the dynamic switching between the typically anti-correlated FPN and DMN, whose coupling is state-dependent and can vary with task demands (V. Menon & D’Esposito, 2022; Tripathi et al., 2025; Uddin et al., 2019). Through this switching, attention is regulated between cognitively demanding, goal-directed behavior on the one hand, and self-referential processing, mentalizing, social cognition, and future planning on the other. Both these processes and networks supporting them are frequently reported to be disrupted in depression (B. Menon, 2019; Prompiengchai & Dunlop, 2025).

Behaviorally, anterior-central clusters were more strongly associated with cognitive control functions such as working memory and inhibitory control as well as the processing of negative emotional stimuli, although at a subthreshold level (**Figure 5** for reference). This functional profile is consistent with the suggested roles of the networks that are connected to the anterior-central clusters: The SAN’s involvement in active detection and prioritization of salient stimuli (V. Menon & Uddin, 2010; Seeley et al., 2007; Sridharan et al., 2008), including emotionally relevant ones. Furthermore, its negative coupling with DMN is consistent with the idea that externally oriented, cognitively demanding tasks such as working memory and inhibitory control require active suppression of internal distractions, functions associated with the DMN (Alves et al., 2019; M. D. Fox et al., 2005; Raichle et al., 2001). The negative connectivity with limbic regions, significantly found for SGC-in the overlap map, would be compatible with a role of anterior-central TMS-PFC in cognitive regulation of emotional responses. Notably, Siddiqi et al. (2021) identified effective TMS and lesion sites converging on a common causal depression circuit anchored in anterolateral DLPFC regions closely overlapping with our anterior-central TMS-PFC subregions and presenting similar connectivity profiles. Our findings refine this picture by revealing substantial functional and behavioral heterogeneity within the anterolateral subregions, particularly between clusters six, two, and eight, that the coarse-level circuit mapping cannot capture and that may be relevant at the spatial scale of millimeter-scale coil placement.

### Posterior areas

In strong contrast to anterior-central clusters, posterior areas, particularly superior-posterior cluster one and the SGC+ region in the overlap map, exhibited the inverse connectivity pattern, with positive RSFC to DMN and negative connectivity to SAN and DAN. These regions showed stronger behavioral associations with explicit memory, social cognition, and positive emotional processing (**Figure 5** for reference). This aligns well with the established DMN functions in explicit (episodic) memory retrieval, mentalizing, theory of mind, and self-referential thought (Alves et al., 2019; M. D. Fox et al., 2005; Raichle et al., 2001). The negative coupling of the superior-posterior cluster with SAN is consistent with internally focused processing, a pattern associated with external stimuli being attenuated to facilitate introspection and memory retrieval. Although the superior-posterior cluster one, the anteriorly adjacent cluster five, and the ventrally adjacent cluster seven shared broadly similar DMN-positive and SAN/DAN-negative connectivity, the cluster-specific conjunction analysis revealed important distinctions: Cluster one alone exhibited the strongest specific connectivity with DMN and anti-correlation with SAN when contrasted against all other clusters, representing the most strongly differentiated posterior pole of this organization. Consistent with Siddiqi et al. (2020), who identified posterior DLPFC and mPFC targets as showing greater efficacy for anxiety and somatic symptoms in depression, the superior-posterior cluster’s network profile, characterized by DMN coupling and SAN anti-correlation, may be particularly relevant when targeting these specific symptom profiles. Clusters five and seven showed additional involvement with somatomotor and visual networks, respectively, suggesting partially distinct functional roles despite their spatial proximity. These partially distinct functional roles show that spatial differentiation within posterior TMS-PFC is as pronounced as in the anterior region, warranting further exploration in future TMS targeting studies.

#### Anterior-central cluster eight

Although the anterior-central cluster eight broadly reflected the triple-network profile of the dorsally adjacent cluster six, its connectivity pattern was not identical. Cluster eight showed cluster-specific positive connectivity with FPN and DAN, particularly with inferior posterior temporal gyrus, SPL, and ventral prefrontal regions, distinguishing it from cluster six, which more specifically engaged SAN. Behaviorally, this cluster was significantly associated with working memory and reasoning and language processes, albeit not reaching significance. Anatomically, its position aligns with the FPN (Yeo et al., 2011), although the FPN spans multiple TMS-PFC regions beyond this cluster. Evidence suggests that the FPN may be fractionated into at least two subnetworks: One coupled with DMN supporting the regulation of introspective processes, and another more connected with DAN and engaged in the regulation of perceptual attention (Corbetta & Shulman, 2002; Uddin et al., 2019). Cluster eight’s profile more closely resembles the latter, which has been linked to voluntary and goal-directed orienting of visuospatial attention (Dixon et al., 2018). These network-level associations are further underscored by reports of hypoconnectivity within DAN (Sacchet et al., 2016; Yu et al., 2019), as well as between FPN and DAN (Kaiser et al., 2015; Sacchet et al., 2016) in depression, although these are less consistently reported than the triple-network disruptions. Together, the distinct profiles of clusters six and eight highlight further functional specificity along the ventral-to-dorsal axis among the anterior subregions.

#### Ventral areas

Complementing the differentiated functional and behavioral profiles of the other clusters, the more ventral clusters (five, seven, and nine) exhibited distinctive connectivity with sensorimotor and visual networks. These clusters were also associated with behavioral domains of visual color and language, e.g., clusters five and nine. Most notably, the most inferior subregion (cluster nine) was primarily linked to language and memory processes, consistent with its anatomical proximity to classical Broca’s area (Broca, 1861; Fedorenko & Blank, 2020), and showed positive connectivity with SMN and negative connectivity with SAN and FPN. Altered SMN function has also been implicated in depression, with studies reporting within-SMN hypoconnectivity (Yan et al., 2019) and decreased regional homogeneity (Iwabuchi et al., 2015), both of which may contribute to psychomotor retardation, a common depressive symptom (Buyukdura et al., 2011). In contrast, the dorsally adjacent clusters five and seven displayed negative connectivity with SMN and VIS. This pattern warrants further exploration in clinical samples, as some studies have reported hypoconnectivity not only within SMN and VIS but also between them (Yan et al., 2019), decreased nodal degree and efficiency across SMN, VIS, DMN, and DAN (Yang et al., 2021), whereas others have found hyperconnectivity within SMN and VIS in major depressive disorder (Yu et al., 2019). Interestingly, cluster five also demonstrated positive connectivity with FPN, whereas cluster nine showed negative connectivity with FPN, suggesting complementary roles at the interface of sensory-motor and executive control functions.

#### Anterior-superior areas

Extending this profile, anterior-superior clusters two and three showed distinct significant associations with social cognition, though with different emphases. The more anterior cluster two followed the general TMS-PFC anterior pattern, showing positive connectivity with SAN and VIS and negative coupling with DMN, LIM, and FPN. It also showed associations with perceptual processes, thermoregulation, and subthreshold involvement in inhibitory control and negative emotions. In contrast, the posteriorly adjacent cluster three aligned more with the posterior pattern, exhibiting positive connectivity with DMN and anticorrelation with FPN and DAN, along with associations with positive emotions and explicit memory, though at a subthreshold level.

#### Areas with constrained cluster-specific network profiles

The superior clusters two and three, as well as the central clusters four and five, exhibited more constrained cluster-specific connectivity compared to the more pronounced antagonistic patterns described above. While spatial autocorrelation of the BOLD signal cannot be fully ruled out as a contributing factor, these clusters nonetheless showed meaningful RSFC and behavioral associations in their main effects (**Figure 4A** for reference). From a neuromodulation perspective, limited cluster-specific network embedding may itself be informative, suggesting that these subregions are less sharply differentiated from their neighbors in terms of the circuits they engage, a distinction that would need to be tested against stimulation outcomes in clinical populations.

These clusters are also among the smallest in our parcellation, particularly clusters three to five (1.13– 2.88 cm^3^, compared with a mean cluster volume of 3.76 cm^3^), raising the question of whether such subdivisions are clinically useful given that activation volumes from electric-field models of standard figure-eight TMS coils range between 3 and 7 cm^3^ (M. D. Fox et al., 2013; Konakanchi et al., 2020; Thielscher et al., 2015). We would argue, however, that this reflects a current constraint on stimulation specificity rather than on the description of TMS-PFC functional organization. Since field intensity decays with distance, targeting the peak-probability voxel still delivers the strongest stimulation to the intended subregion. Nonetheless, co-stimulation of adjacent subregions with engagement of distributed network effects, although of lesser magnitude, would be expected, suggesting these smaller clusters as less precise targets when the aim is to engage specific functional circuits.

Taken together, these findings highlight that while many TMS-PFC clusters followed an antagonistic triple-network pattern (DMN-SAN-FPN), others diverged toward more specialized roles. For instance, the involvement of central clusters four and five in color processing underscores integration with perceptual systems, whereas cluster nine’s language-related profile and SMN connectivity suggest potential links to psychomotor and communication processes. Such heterogeneity indicates that not all TMS-PFC subregions can be reduced to the dominant anterior-posterior antagonism but instead support distinct functions. Whether these specialized subregions are relevant for targeting specific symptom dimensions, such as language deficits, perceptual disturbances, or psychomotor symptoms, remains an open question for future clinical studies. In summary, our results reveal a strongly antagonistic organization along the anterior-central to superior-posterior axis of the TMS-PFC, alongside distinct ventral and anterior-superior clusters with stronger involvement in social cognition, sensorimotor integration, and language-related processes. This layered organization underscores the functional heterogeneity of TMS-PFC and provides a reference from which hypotheses about individualized and symptom-specific stimulation strategies can be derived and tested.

### 4.2 Overlap Map and Clinical Implications

Building on the fine-grained parcellation, we derived an overlap map of TMS-PFC integrating SGC connectivity patterns across multiple granularities (**Figure 6A** for reference). Unlike a direct voxel-wise SGC connectivity map, which reflects average connectivity at a single level of analysis, the overlap map captures the spatial consistency of antagonistic SGC connectivity patterns across both biologically plausible granularities and individuals. This provides a more robust reference than any single connectivity threshold or parcellation solution. Importantly, the overlap map is not intended to replace the nine-cluster solution but to complement it: While the nine-cluster solution characterizes the full functional heterogeneity of TMS-PFC, the overlap map provides a tractable summary of SGC connectivity that situates our results within the well-established SGC-centric framework of TMS, which links stronger SGC anti-correlation of the stimulation site to better antidepressant outcomes (Cash, Cocchi, Lv, Fitzgerald, et al., 2021; Cash et al., 2019; M. D. Fox et al., 2012, 2013; Siddiqi et al., 2021; Weigand et al., 2018). Beyond SGC connectivity, the map also captures the antagonistic triple-network pattern between DMN, SAN, and FPN, relating our findings to circuit-level models for depression.

From a clinical perspective, the functional organization of the TMS-PFC subregions derived from our clustering may serve as a reference for future studies analyzing dysregulations in depression. In particular, the anterior-central SGC-region, consistently anti-correlated with SGC, DMN and LIM and positively connected with SAN, FPN, and DAN, was associated with working memory, action inhibition, sadness, and pain perception, functions commonly affected in depression (Fried & Nesse, 2015; Northoff & Hirjak, 2024). In contrast, the superior-posterior SGC+ region, positively correlated with SGC and DMN and negatively with SAN and DAN, showed stronger associations with explicit memory, social cognition, and positive emotional processes, domains often reported as disrupted in depression in the context of anhedonia and social withdrawal (Fried & Nesse, 2015; Northoff & Hirjak, 2024). These findings resemble the dual-circuit framework proposed by Cash et al. (2023), distinguishing separate emotion- and cognition-related networks involving the DLPFC in depression, and are broadly consistent with the symptom-circuit mapping of Siddiqi et al. (2020), linking anterolateral DLPFC-SGC sites to dysphoric symptoms and posterior targets to anxiety and somatic complaints.

Functionally, the anterior-central SGC-shows a connectivity profile consistent with a regulatory role in cognitive and emotional control, whereas the superior-posterior SGC+ may be more involved in internally directed, emotionally valenced cognition. Given reports of altered connectivity in depressive disorder, most notably increased intra-network DMN connectivity and reduced anti-correlation with attentional and control networks (Scalabrini et al., 2020; Tripathi et al., 2025), and the link between DMN functioning, rumination, and self-referential processing in depression (Northoff & Hirjak, 2024; Sheline et al., 2009), both the anterior-central SGC- and superior-posterior SGC+ emerge as potential stimulation targets warranting investigation in clinical populations. In line with the triple-network model of psychopathology (B. Menon, 2019; V. Menon, 2011), the anterior-central SGC-may be of particular interest for modulating dysfunctional large-scale interactions. Should stimulation of this region influence the reduced anti-correlation between DMN and attentional/control networks, it may facilitate the recovery of functional segregation between self-referential and executive processes, potentially ameliorating symptoms such as biased attentional processing and impaired inhibition of ruminative thoughts into negative emotional content (Fossati, 2019; Javaheripour et al., 2023; Lynch et al., 2024; Northoff & Hirjak, 2024). This remains a hypothesis to be tested against treatment outcomes.

At the same time, our findings caution against a one-size-fits-all perspective. Depression is a heterogeneous disorder, and while the anterior-central SGC- and superior-posterior SGC+ regions may be most relevant for cognitive-affective symptoms, other clusters, with their distinct sensorimotor, language, and social cognition profiles, may be more pertinent for motor, perceptual, or interpersonal disturbances. This is compatible with emerging symptom-guided neuromodulation strategies (Siddiqi et al., 2020) and underscores the substantial inter-individual variability in the precise location and extent of SGC connectivity. This highlights the need for individualized targeting approaches that balance tractable group-level heuristics with patient-specific connectivity profiles. Our parcellation is defined at the group level, but it may serve as a spatial prior for such approaches, constraining the search space for individual optimization within functionally defined subregions.

All in all, our fine-grained parcellation and overlap map extend the SGC-centric framework of TMS targeting by identifying SGC-as a leading candidate for future targeting studies, while simultaneously demonstrating that adjacent subregions differ markedly in network embedding and behavioral relevance. This suggests that therapeutic benefit may depend less on stimulating a fixed coordinate than on modulating distributed large-scale networks whose balance varies across individuals (Moreno-Ortega et al., 2020).

### 4.3 Limitations and Future Outlook

A key limitation of this work is that our analysis was conducted on the HCP-YA dataset, which consists of young, healthy individuals scanned with high-quality acquisition protocols. While this ensured high signal-to-noise ratios and reliable connectivity estimates, it limits the generalizability of our findings to more diverse and clinically relevant populations, whose data are typically noisier. It remains to be established whether the functional organization we observed between TMS-PFC subregions in healthy individuals is preserved or altered in patients with TRD. Relatedly, our study includes no TMS treatment data or clinical outcome measures. Future research should therefore examine these subregions in clinical populations before and after treatment, to identify potential biomarkers of treatment response or maladaptive network dynamics underlying specific depressive symptom dimensions. Within this broader framework, the anterior-central subregion is therefore only presented here as a candidate target that still requires clinical validation.

In addition, the selection of an optimal clustering solution in connectivity-based parcellation remains challenging, as no single criterion can unambiguously identify the “true” number of neurobiologically meaningful subregions. Our heuristic convergence approach across multiple criteria mitigates but does not fully resolve this limitation. Relatedly, we did not employ a null model based on random parcellations (e.g., permutation-based comparison against random rCBP solutions) as a formal selection criterion. However, such approaches address statistical significance rather than functional optimality, and generating hundreds of individual-level random parcellations for a large dataset and nine levels of granularity is computationally highly demanding.

A further consideration concerns the spatial resolution of the smaller subregions in our nine-cluster parcellation relative to the broad electric field (e-field) induced by TMS. Our aim was to provide a reference that constrains the search space for target optimization, such that stimulation can be centered where the intended functional area is engaged with the highest probability. As discussed above, the induced e-field is nevertheless broad, and its distribution depends on coil orientation, scalp-to-cortex distance, cortical folding, and stimulation intensity, such that neighboring subregions separated by only a few millimeters may be co-stimulated. These physical TMS parameters were not modeled here but warrant attention in future work combining parcellation-based targeting with e-field modeling.

Moreover, we see a pressing need for greater standardization and transparency in TMS research. We strongly advocate for future publications to report not only their stimulation coordinates but also the specific MNI space and transformation procedures used and, where possible, clinical outcome scores. Such practices would greatly enhance reproducibility, allow for precise cross-study comparisons, and ultimately enable meta-analyses that could accelerate the development of clinically relevant treatment pipelines. Such standardization would also help clarify whether observed heterogeneity in clinical outcomes reflects true neurobiological differences or methodological inconsistencies in target localization.

Building on our findings, an important next step is to bridge the group-level organization described here with individual-level targeting. Individual connectivity estimates are considerably noisy, and single-voxel targeting is correspondingly susceptible to measurement noise. Thus, a group-level parcellation may help by constraining the search space for the optimized selection of individualized stimulation targets: Rather than searching the entire prefrontal cortex or relying on single-voxel connectivity maps, stimulation sites could be optimized within functionally defined subregions. For example, with a particular focus on the anterior-central TMS-PFC subregion to maximize anti-correlation with SGC and DMN while concurrently enhancing connectivity with SAN, DAN, and control networks. Such an approach could extend the current strategy of targeting the most SGC-anticorrelated site toward models that also account for broader network dynamics (Moreno-Ortega et al., 2020), though its feasibility and clinical value remain to be tested. Importantly, our results also raise questions about the role of the superior-posterior cluster, which shows a pattern associated with positive emotional processing, social cognition, and positive DMN connectivity. Rather than ignoring them, such regions might be examined in future studies to determine whether they should be actively avoided due to potentially counterproductive effects or, alternatively, even targeted in specific depressive subtypes (Cash et al., 2023; Duprat et al., 2025; Oathes et al., 2023; Siddiqi et al., 2020).

## 5 Conclusion

Our findings demonstrate that the left prefrontal cortex, within a clinically relevant stimulation region of interest, contains multiple functionally distinct subregions that differ in their connectivity with large-scale functional networks, as well as in their associated behavioral profiles. This organization can be captured at two complementary hierarchical levels: A coarse two-pole antagonism (anterior-central vs. superior-posterior) and a finer nine-cluster architecture that reveals the TMS-PFC’s functional heterogeneity in greater detail along anterior-to-posterior as well as ventral-to-dorsal axes. By integrating these perspectives, our work refines the understanding of prefrontal functional organization and provides a network-informed TMS-PFC functional atlas that can inform more precise and individualized TMS targeting strategies, pending clinical validation. Future work should examine the relevance of these subdivisions in patients with depression and their potential to optimize treatment outcomes.

## Data and Code Availability

This study used openly available data from the Human Connectome Project Young Adult dataset (https://www.humanconnectome.org/study/hcp-young-adult). The code for the analyses reported in this manuscript is available on GitHub (https://github.com/lyapaas/ldlpfc_parcellation). The source code for the connectivity-based parcellation used in this manuscript is openly available at https://github.com/juaml/rcbp, which is an adaptation from https://github.com/inm7/cbptools/ for the specific computation demands. The documentation for CBPtools (Reuter et al., 2020) can be found at https://cbptools.readthedocs.io/.

## Author Contributions

**LKPO:** Conceptualization, Software, Formal analysis, Investigation, Visualization, Writing - Original Draft, Writing - Review & Editing. **TBP:** Conceptualization, Investigation, Methodology, Writing - Review & Editing, Funding acquisition. **NR:** Methodology, Software. **KRP:** Conceptualization, Methodology. **SK:** Validation, Writing - Review & Editing. **NYT:** Validation, Writing - Review & Editing. **RFHC:** Validation, Writing - Review & Editing. **FH:** Validation, Writing - Review & Editing. **SBE:** Conceptualization, Investigation, Methodology, Resources, Writing - Review & Editing, Supervision, Funding acquisition. **VIM:** Conceptualization, Investigation, Methodology, Writing - Review & Editing, Supervision.

## Acknowledgments

A version of this article was published as a preprint on bioRxiv: https://doi.org/10.1101/2025.11.19.689337. Data were provided by the Human Connectome Project, WU-Minn Consortium (Principal Investigators: David Van Essen and Kamil Ugurbil; 1U54MH091657) funded by the 16 NIH Institutes and Centers that support the NIH Blueprint for Neuroscience Research; and by the McDonnell Center for Systems Neuroscience at Washington University, USA. Open Access funding enabled and organized by Projekt DEAL.

## Declaration of generative AI use

During the preparation of this work, ChatGPT and Claude were used for language editing and improving readability. After using these tools, the author reviewed and edited the content as needed and takes full responsibility for the content of the published article.

## References

Abdallah, M., Zanitti, G. E., Iovene, V., & Wassermann, D. (2022). Functional gradients in the human lateral prefrontal cortex revealed by a comprehensive coordinate-based meta-analysis. eLife, 11, e76926. 10.7554/elife.76926

Alexander-Bloch, A. F., Shou, H., Liu, S., Satterthwaite, T. D., Glahn, D. C., Shinohara, R. T., Vandekar, S. N., & Raznahan, A. (2018). On testing for spatial correspondence between maps of human brain structure and function. NeuroImage, 178, 540–551. 10.1016/j.neuroimage.2018.05.070

Alves, P. N., Foulon, C., Karolis, V., Bzdok, D., Margulies, D. S., Volle, E., & Schotten, M. T. de. (2019). An improved neuroanatomical model of the default-mode network reconciles previous neuroimaging and neuropathological findings. Communications Biology, 2(1), 370. 10.1038/s42003-019-0611-3

Amunts, K., Mohlberg, H., Bludau, S., & Zilles, K. (2020). Julich-Brain: A 3D probabilistic atlas of the human brain’s cytoarchitecture. Science, 369(6506), 988–992. 10.1126/science.abb4588

Amunts, K., Mohlberg, H., Buldau, S., Caspers, S., Lewis, L. B., Eickhoff, S. B., & Pieperhoff, P. (2023). Julich-Brain Atlas, cytoarchitectonic maps (v3.1) [Data set].

Anderson, R. J., Hoy, K. E., Daskalakis, Z. J., & Fitzgerald, P. B. (2016). Repetitive transcranial magnetic stimulation for treatment resistant depression: Re-establishing connections. Clinical Neurophysiology, 127(11), 3394–3405. 10.1016/j.clinph.2016.08.015

Arslan, S., Ktena, S. I., Makropoulos, A., Robinson, E. C., Rueckert, D., & Parisot, S. (2018). Human brain mapping: A systematic comparison of parcellation methods for the human cerebral cortex. NeuroImage, 170, 5–30. 10.1016/j.neuroimage.2017.04.014

Beckmann, C. F., & Smith, S. M. (2004). Probabilistic Independent Component Analysis for Functional Magnetic Resonance Imaging. IEEE Transactions on Medical Imaging, 23(2), 137–152. 10.1109/tmi.2003.822821

Beynel, L., Powers, J. P., & Appelbaum, L. G. (2020). Effects of repetitive transcranial magnetic stimulation on resting-state connectivity: A systematic review. NeuroImage, 211, 116596. 10.1016/j.neuroimage.2020.116596

Bijsterbosch, J., Harrison, S. J., Jbabdi, S., Woolrich, M., Beckmann, C., Smith, S., & Duff, E. P. (2020). Challenges and future directions for representations of functional brain organization. Nature Neuroscience, 23(12), 1484–1495. 10.1038/s41593-020-00726-z

Blumberger, D. M., Vila-Rodriguez, F., Thorpe, K. E., Feffer, K., Noda, Y., Giacobbe, P., Knyahnytska, Y., Kennedy, S. H., Lam, R. W., Daskalakis, Z. J., & Downar, J. (2018). Effectiveness of theta burst versus high-frequency repetitive transcranial magnetic stimulation in patients with depression (THREE-D): a randomised non-inferiority trial. The Lancet, 391(10131), 1683–1692. 10.1016/s0140-6736(18)30295-2

Broca, P. (1861). Remarques Sur le Siége de la Faculté Du Langage Articulé, Suivies D’une Observation D’aphémie (Perte de la Parole). Bulletin Society Anatomique, (6), 330–357.

Brodmann, K. (1909). *Vergleichende Lokalisationslehre der Grosshirnrinde in ihren Prinzipien dargestellt auf Grund des Zellenbaues* (L. J. A. Barth, Ed.).

Bruno, A., Lothmann, K., Bludau, S., Mohlberg, H., & Amunts, K. (2024). New organizational principles and 3D cytoarchitectonic maps of the dorsolateral prefrontal cortex in the human brain. Frontiers in Neuroimaging, 3, 1339244. 10.3389/fnimg.2024.1339244

Buyukdura, J. S., McClintock, S. M., & Croarkin, P. E. (2011). Psychomotor retardation in depression: Biological underpinnings, measurement, and treatment. Progress in Neuro-Psychopharmacology and Biological Psychiatry, 35(2), 395–409. 10.1016/j.pnpbp.2010.10.019

Cash, R. F. H., Cocchi, L., Lv, J., Fitzgerald, P. B., & Zalesky, A. (2021). Functional Magnetic Resonance Imaging– Guided Personalization of Transcranial Magnetic Stimulation Treatment for Depression. JAMA Psychiatry, 78(3), 337–339. 10.1001/jamapsychiatry.2020.3794

Cash, R. F. H., Cocchi, L., Lv, J., Wu, Y., Fitzgerald, P. B., & Zalesky, A. (2021). Personalized connectivity-guided DLPFC-TMS for depression: Advancing computational feasibility, precision and reproducibility. Human Brain Mapping, 42(13), 4155–4172. 10.1002/hbm.25330

Cash, R. F. H., Müller, V. I., Fitzgerald, P. B., Eickhoff, S. B., & Zalesky, A. (2023). Altered brain activity in unipolar depression unveiled using connectomics. Nature Mental Health, 1(3), 174–185. 10.1038/s44220-023-00038-8

Cash, R. F. H., Weigand, A., Zalesky, A., Siddiqi, S. H., Downar, J., Fitzgerald, P. B., & Fox, M. D. (2021). Using Brain Imaging to Improve Spatial Targeting of Transcranial Magnetic Stimulation for Depression. Biological Psychiatry, 90(10), 689–700. 10.1016/j.biopsych.2020.05.033

Cash, R. F. H., & Zalesky, A. (2024). Personalized and Circuit-Based Transcranial Magnetic Stimulation: Evidence, Controversies, and Opportunities. Biological Psychiatry, 95(6), 510–522. 10.1016/j.biopsych.2023.11.013

Cash, R. F. H., Zalesky, A., Thomson, R. H., Tian, Y., Cocchi, L., & Fitzgerald, P. B. (2019). Subgenual Functional Connectivity Predicts Antidepressant Treatment Response to Transcranial Magnetic Stimulation: Independent Validation and Evaluation of Personalization. Biological Psychiatry, 86(2), e5–e7. 10.1016/j.biopsych.2018.12.002

Castrillon, G., Sollmann, N., Kurcyus, K., Razi, A., Krieg, S. M., & Riedl, V. (2020). The physiological effects of noninvasive brain stimulation fundamentally differ across the human cortex. Science Advances, 6(5), eaay2739. 10.1126/sciadv.aay2739

Chai, Y., Sheline, Y. I., Oathes, D. J., Balderston, N. L., Rao, H., & Yu, M. (2023). Functional connectomics in depression: insights into therapies. Trends in Cognitive Sciences, 27(9), 814–832. 10.1016/j.tics.2023.05.006

Cieslik, E. C., Zilles, K., Caspers, S., Roski, C., Kellermann, T. S., Jakobs, O., Langner, R., Laird, A. R., Fox, P. T., & Eickhoff, S. B. (2013). Is There “One” DLPFC in Cognitive Action Control? Evidence for Heterogeneity From Co-Activation-Based Parcellation. Cerebral Cortex, 23(11), 2677–2689. 10.1093/cercor/bhs256

Cole, E. J., Phillips, A. L., Bentzley, B. S., Stimpson, K. H., Nejad, R., Barmak, F., Veerapal, C., Khan, N., Cherian, K., Felber, E., Brown, R., Choi, E., King, S., Pankow, H., Bishop, J. H., Azeez, A., Coetzee, J., Rapier, R., Odenwald, N.,…Williams, N. R. (2022). Stanford Neuromodulation Therapy (SNT): A Double-Blind Randomized Controlled Trial. American Journal of Psychiatry, 179(2), 132–141. 10.1176/appi.ajp.2021.20101429

Corbetta, M., & Shulman, G. L. (2002). Control of goal-directed and stimulus-driven attention in the brain. Nature Reviews Neuroscience, 3(3), 201–215. 10.1038/nrn755

Desikan, R. S., Ségonne, F., Fischl, B., Quinn, B. T., Dickerson, B. C., Blacker, D., Buckner, R. L., Dale, A. M., Maguire, R. P., Hyman, B. T., Albert, M. S., & Killiany, R. J. (2006). An automated labeling system for subdividing the human cerebral cortex on MRI scans into gyral based regions of interest. NeuroImage, 31(3), 968–980. 10.1016/j.neuroimage.2006.01.021

Dixon, M. L., Vega, A. D. L., Mills, C., Andrews-Hanna, J., Spreng, R. N., Cole, M. W., & Christoff, K. (2018). Heterogeneity within the frontoparietal control network and its relationship to the default and dorsal attention networks. Proceedings of the National Academy of Sciences, 115(7), E1598–E1607. 10.1073/pnas.1715766115

Doucet, G. E., Lee, W. H., & Frangou, S. (2019). Evaluation of the spatial variability in the major resting-state networks across human brain functional atlases. Human Brain Mapping, 40(15), 4577–4587. 10.1002/hbm.24722

Drevets, W. C., Savitz, J., & Trimble, M. (2008). The Subgenual Anterior Cingulate Cortex in Mood Disorders. CNS Spectrums, 13(8), 663–681. 10.1017/s1092852900013754

Drysdale, A. T., Grosenick, L., Downar, J., Dunlop, K., Mansouri, F., Meng, Y., Fetcho, R. N., Zebley, B., Oathes, D. J., Etkin, A., Schatzberg, A. F., Sudheimer, K., Keller, J., Mayberg, H. S., Gunning, F. M., Alexopoulos, G. S., Fox, M. D., Pascual-Leone, A., Voss, H. U.,…Liston, C. (2017). Resting-state connectivity biomarkers define neurophysiological subtypes of depression. Nature Medicine, 23(1), 28–38. 10.1038/nm.4246

Duprat, R. J., Linn, K. A., Satterthwaite, T. D., Sheline, Y. I., Liang, X., Bagdon, G., Flounders, M. W., Robinson, H., Platt, M., Kable, J., Long, H., Scully, M., Deluisi, J. A., Thase, M., Cristancho, M., Grier, J., Blaine, C., Figueroa-González, A., & Oathes, D. J. (2025). Resting fMRI-guided TMS evokes subgenual anterior cingulate response in depression. NeuroImage, 305, 120963. 10.1016/j.neuroimage.2024.120963

Eickhoff, S. B., Thirion, B., Varoquaux, G., & Bzdok, D. (2015). Connectivity-based parcellation: Critique and implications. Human Brain Mapping, 36(12), 4771–4792. 10.1002/hbm.22933

Eickhoff, S. B., Yeo, B. T. T., & Genon, S. (2018). Imaging-based parcellations of the human brain. Nature Reviews Neuroscience, 19(11), 672–686. 10.1038/s41583-018-0071-7

European Medicines Agency. (2025). *Guideline on clinical investigation of medicinal products in the treatment of depression*. European Medicines Agency. https://www.ema.europa.eu/en/documents/scientific-guideline/guideline-clinical-investigation-medicinal-products-treatment-depression-revision-3_en.pdf

Fedorenko, E., & Blank, I. A. (2020). Broca’s Area Is Not a Natural Kind. Trends in Cognitive Sciences, 24(4), 270– 284. 10.1016/j.tics.2020.01.001

Fitzgerald, P. B., Hoy, K. E., Anderson, R. J., & Daskalakis, Z. J. (2016). A study of the pattern of response to rTMS treatment in depression. Depression and Anxiety, 33(8), 746–753. 10.1002/da.22503

Fitzgerald, P. B., Laird, A. R., Maller, J., & Daskalakis, Z. J. (2008). A meta-analytic study of changes in brain activation in depression. Human Brain Mapping, 29(6), 683–695. 10.1002/hbm.20426

Fossati, P. (2019). Circuit based anti-correlation, attention orienting, and major depression. CNS Spectrums, 24(1), 94–101. 10.1017/s1092852918001402

Fox, M. D., Buckner, R. L., White, M. P., Greicius, M. D., & Pascual-Leone, A. (2012). Efficacy of Transcranial Magnetic Stimulation Targets for Depression Is Related to Intrinsic Functional Connectivity with the Subgenual Cingulate. Biological Psychiatry, 72(7), 595–603. 10.1016/j.biopsych.2012.04.028

Fox, M. D., Liu, H., & Pascual-Leone, A. (2013). Identification of reproducible individualized targets for treatment of depression with TMS based on intrinsic connectivity. NeuroImage, 66, 151–160. 10.1016/j.neuroimage.2012.10.082

Fox, M. D., Snyder, A. Z., Vincent, J. L., Corbetta, M., Essen, D. C. V., & Raichle, M. E. (2005). The human brain is intrinsically organized into dynamic, anticorrelated functional networks. Proceedings of the National Academy of Sciences, 102(27), 9673–9678. 10.1073/pnas.0504136102

Fox, P. T., & Lancaster, J. L. (2002). Mapping context and content: the BrainMap model. Nature Reviews Neuroscience, 3(4), 319–321. 10.1038/nrn789

Fried, E. I., & Nesse, R. M. (2015). Depression sum-scores don’t add up: why analyzing specific depression symptoms is essential. BMC Medicine, 13(1), 72. 10.1186/s12916-015-0325-4

Friedman, N. P., & Miyake, A. (2017). Unity and diversity of executive functions: Individual differences as a window on cognitive structure. Cortex, 86, 186–204. 10.1016/j.cortex.2016.04.023

Gale, D. J., Wael, R. V. de, Benkarim, O., & Bernhardt, B. (2021). Surfplot: Publication-ready brain surface figures (v0.1.0). Zenodo. 10.5281/zenodo.5567926

Genon, S., Li, H., Fan, L., Müller, V. I., Cieslik, E. C., Hoffstaedter, F., Reid, A. T., Langner, R., Grefkes, C., Fox, P. T., Moebus, S., Caspers, S., Amunts, K., Jiang, T., & Eickhoff, S. B. (2017). The Right Dorsal Premotor Mosaic: Organization, Functions, and Connectivity. Cerebral Cortex, 27(3), 2095–2110. 10.1093/cercor/bhw065

Genon, S., Reid, A., Langner, R., Amunts, K., & Eickhoff, S. B. (2018). How to Characterize the Function of a Brain Region. Trends in Cognitive Sciences, 22(4), 350–364. 10.1016/j.tics.2018.01.010

Glasser, M. F., Sotiropoulos, S. N., Wilson, J. A., Coalson, T. S., Fischl, B., Andersson, J. L., Xu, J., Jbabdi, S., Webster, M., Polimeni, J. R., Essen, D. C. V., Jenkinson, M., & for the WU-Minn HCP Consortium. (2013). The minimal preprocessing pipelines for the Human Connectome Project. NeuroImage, 80, 105–124. 10.1016/j.neuroimage.2013.04.127

Godfrey, K. E. M., Muthukumaraswamy, S. D., Stinear, C. M., & Hoeh, N. (2022). Decreased salience network fMRI functional connectivity following a course of rTMS for treatment-resistant depression. Journal of Affective Disorders, 300, 235–242. 10.1016/j.jad.2021.12.129

Gratton, C., Lee, T. G., Nomura, E. M., & D’Esposito, M. (2013). The effect of theta-burst TMS on cognitive control networks measured with resting state fMRI. Frontiers in Systems Neuroscience, 7, 124. 10.3389/fnsys.2013.00124

Griffanti, L., Salimi-Khorshidi, G., Beckmann, C. F., Auerbach, E. J., Douaud, G., Sexton, C. E., Zsoldos, E., Ebmeier, K. P., Filippini, N., Mackay, C. E., Moeller, S., Xu, J., Yacoub, E., Baselli, G., Ugurbil, K., Miller, K. L., & Smith, S. M. (2014). ICA-based artefact removal and accelerated fMRI acquisition for improved resting state network imaging. NeuroImage, 95, 232–247. 10.1016/j.neuroimage.2014.03.034

Herbsman, T., Avery, D., Ramsey, D., Holtzheimer, P., Wadjik, C., Hardaway, F., Haynor, D., George, M. S., & Nahas, Z. (2009). More Lateral and Anterior Prefrontal Coil Location Is Associated with Better Repetitive Transcranial Magnetic Stimulation Antidepressant Response. Biological Psychiatry, 66(5), 509–515. 10.1016/j.biopsych.2009.04.034

Institute for Health Metrics and Evaluation. (2020). GBD Results. Institute for Health Metrics and Evaluation, University of Washington. https://vizhub.healthdata.org/gbd-results/

Iwabuchi, S. J., Krishnadas, R., Li, C., Auer, D. P., Radua, J., & Palaniyappan, L. (2015). Localized connectivity in depression: A meta-analysis of resting state functional imaging studies. Neuroscience & Biobehavioral Reviews, 51, 77–86. 10.1016/j.neubiorev.2015.01.006

Javaheripour, N., Colic, L., Opel, N., Li, M., Balajoo, S. M., Chand, T., Meer, J. V. der, Krylova, M., Izyurov, I., Meller, T., Goltermann, J., Winter, N. R., Meinert, S., Grotegerd, D., Jansen, A., Alexander, N., Usemann, P., Thomas-Odenthal, F., Evermann, U.,…Walter, M. (2023). Altered brain dynamic in major depressive disorder: state and trait features. Translational Psychiatry, 13(1), 261. 10.1038/s41398-023-02540-0

Jung, J., Ralph, M. A. L., & Jackson, R. L. (2022). Subregions of DLPFC Display Graded yet Distinct Structural and Functional Connectivity. The Journal of Neuroscience, 42(15), 3241–3252. 10.1523/jneurosci.1216-21.2022

Kaiser, R. H., Andrews-Hanna, J. R., Wager, T. D., & Pizzagalli, D. A. (2015). Large-Scale Network Dysfunction in Major Depressive Disorder: A Meta-analysis of Resting-State Functional Connectivity. JAMA Psychiatry, 72(6), 603–611. 10.1001/jamapsychiatry.2015.0071

Koenigs, M., & Grafman, J. (2009). The functional neuroanatomy of depression: Distinct roles for ventromedial and dorsolateral prefrontal cortex. Behavioural Brain Research, 201(2), 239–243. 10.1016/j.bbr.2009.03.004

Konakanchi, D., Curry, A. L. de J., Waters, R. S., & Narayana, S. (2020). Focality of the Induced E-Field Is a Contributing Factor in the Choice of TMS Parameters: Evidence from a 3D Computational Model of the Human Brain. Brain Sciences, 10(12), 1010. 10.3390/brainsci10121010

Kong, R., Spreng, R. N., Xue, A., Betzel, R. F., Cohen, J. R., Damoiseaux, J. S., Brigard, F. D., Eickhoff, S. B., Fornito, A., Gratton, C., Gordon, E. M., Holmes, A. J., Laird, A. R., Larson-Prior, L., Nickerson, L. D., Pinho, A. L., Razi, A., Sadaghiani, S., Shine, J. M.,…Uddin, L. Q. (2025). A network correspondence toolbox for quantitative evaluation of novel neuroimaging results. Nature Communications, 16(1), 2930. 10.1038/s41467-025-58176-9

Kraskov, A., Stögbauer, H., Andrzejak, R. G., & Grassberger, P. (2005). Hierarchical clustering using mutual information. EPL (Europhysics Letters), 70(2), 278–284. 10.1209/epl/i2004-10483-y

Laird, A. R., Eickhoff, S. B., Fox, P. M., Uecker, A. M., Ray, K. L., Saenz, J. J., McKay, D. R., Bzdok, D., Laird, R. W., Robinson, J. L., Turner, J. A., Turkeltaub, P. E., Lancaster, J. L., & Fox, P. T. (2011). The BrainMap strategy for standardization, sharing, and meta-analysis of neuroimaging data. BMC Research Notes, 4(1), 349. 10.1186/1756-0500-4-349

Laird, A. R., Lancaster, J. J., & Fox, P. T. (2005). BrainMap: The Social Evolution of a Human Brain Mapping Database. Neuroinformatics, 3(1), 65–77. 10.1385/ni:3:1:065

Li, Y., Li, R., Gu, J., Yi, H., He, J., Lu, F., & Gao, J. (2024). Enhanced group-level dorsolateral prefrontal cortex subregion parcellation through functional connectivity-based distance-constrained spectral clustering with application to autism spectrum disorder. Cerebral Cortex, 34(2), bhae020. 10.1093/cercor/bhae020

Liu, X., Eickhoff, S. B., Caspers, S., Wu, J., Genon, S., Hoffstaedter, F., Mars, R. B., Sommer, I. E., Eickhoff, C. R., Chen, J., Jardri, R., Reetz, K., Dogan, I., Aleman, A., Kogler, L., Gruber, O., Caspers, J., Mathys, C., & Patil, K. R. (2021). Functional parcellation of human and macaque striatum reveals human-specific connectivity in the dorsal caudate. NeuroImage, 235, 118006. 10.1016/j.neuroimage.2021.118006

Liu, X., Eickhoff, S. B., Hoffstaedter, F., Genon, S., Caspers, S., Reetz, K., Dogan, I., Eickhoff, C. R., Chen, J., Caspers, J., Reuter, N., Mathys, C., Aleman, A., Jardri, R., Riedl, V., Sommer, I. E., & Patil, K. R. (2020). Joint Multi-modal Parcellation of the Human Striatum: Functions and Clinical Relevance. Neuroscience Bulletin, 36(10), 1123–1136. 10.1007/s12264-020-00543-1

Lynch, C. J., Elbau, I. G., Ng, T., Ayaz, A., Zhu, S., Wolk, D., Manfredi, N., Johnson, M., Chang, M., Chou, J., Summerville, I., Ho, C., Lueckel, M., Bukhari, H., Buchanan, D., Victoria, L. W., Solomonov, N., Goldwaser, E., Moia, S.,…Liston, C. (2024). Frontostriatal salience network expansion in individuals in depression. Nature, 633(8030), 624–633. 10.1038/s41586-024-07805-2

Mayberg, H. S., Lozano, A. M., Voon, V., McNeely, H. E., Seminowicz, D., Hamani, C., Schwalb, J. M., & Kennedy, S. H. (2005). Deep Brain Stimulation for Treatment-Resistant Depression. Neuron, 45(5), 651–660. 10.1016/j.neuron.2005.02.014

Menon, B. (2019). Towards a new model of understanding – The triple network, psychopathology and the structure of the mind. Medical Hypotheses, 133, 109385. 10.1016/j.mehy.2019.109385

Menon, V. (2011). Large-scale brain networks and psychopathology: a unifying triple network model. Trends in Cognitive Sciences, 15(10), 483–506. 10.1016/j.tics.2011.08.003

Menon, V., & D’Esposito, M. (2022). The role of PFC networks in cognitive control and executive function. Neuropsychopharmacology, 47(1), 90–103. 10.1038/s41386-021-01152-w

Menon, V., & Uddin, L. Q. (2010). Saliency, switching, attention and control: a network model of insula function. Brain Structure and Function, 214(5–6), 655–667. 10.1007/s00429-010-0262-0

Moreno-Ortega, M., Kangarlu, A., Lee, S., Perera, T., Kangarlu, J., Palomo, T., Glasser, M. F., & Javitt, D. C. (2020). Parcel-guided rTMS for depression. Translational Psychiatry, 10(1), 283. 10.1038/s41398-020-00970-8

Mueller, S., Wang, D., Fox, M. D., Yeo, B. T. T., Sepulcre, J., Sabuncu, M. R., Shafee, R., Lu, J., & Liu, H. (2013). Individual Variability in Functional Connectivity Architecture of the Human Brain. Neuron, 77(3), 586– 595. 10.1016/j.neuron.2012.12.028

Nanetti, L., Cerliani, L., Gazzola, V., Renken, R., & Keysers, C. (2009). Group analyses of connectivity-based cortical parcellation using repeated k-means clustering. NeuroImage, 47(4), 1666–1677. 10.1016/j.neuroimage.2009.06.014

Nguyen, N., & Caruana, R. (2007). Consensus Clusterings. Seventh IEEE International Conference on Data Mining, 607–612. 10.1109/icdm.2007.73

Ning, L., Makris, N., Camprodon, J. A., & Rathi, Y. (2019). Limits and reproducibility of resting-state functional MRI definition of DLPFC targets for neuromodulation. Brain Stimulation, 12(1), 129–138. 10.1016/j.brs.2018.10.004

Northoff, G., & Hirjak, D. (2024). Is depression a global brain disorder with topographic dynamic reorganization? Translational Psychiatry, 14(1), 278. 10.1038/s41398-024-02995-9

Oathes, D. J., Duprat, R. J.-P., Reber, J., Liang, X., Scully, M., Long, H., Deluisi, J. A., Sheline, Y. I., & Linn, K. A. (2023). Non-invasively targeting, probing and modulating a deep brain circuit for depression alleviation. Nature Mental Health, 1(12), 1033–1042. 10.1038/s44220-023-00165-2

Palomero-Gallagher, N., Eickhoff, S. B., Hoffstaedter, F., Schleicher, A., Mohlberg, H., Vogt, B. A., Amunts, K., & Zilles, K. (2015). Functional organization of human subgenual cortical areas: Relationship between architectonical segregation and connectional heterogeneity. NeuroImage, 115, 177–190. 10.1016/j.neuroimage.2015.04.053

Panikratova, Y. R., Vlasova, R. M., Akhutina, T. V., Korneev, A. A., Sinitsyn, V. E., & Pechenkova, E. V. (2020). Functional connectivity of the dorsolateral prefrontal cortex contributes to different components of executive functions. International Journal of Psychophysiology, 151, 70–79. 10.1016/j.ijpsycho.2020.02.013

Petrides, M., & Pandya, D. N. (1999). Dorsolateral prefrontal cortex: comparative cytoarchitectonic analysis in the human and the macaque brain and corticocortical connection patterns. European Journal of Neuroscience, 11(3), 1011–1036. 10.1046/j.1460-9568.1999.00518.x

Plachti, A., Eickhoff, S. B., Hoffstaedter, F., Patil, K. R., Laird, A. R., Fox, P. T., Amunts, K., & Genon, S. (2019). Multimodal Parcellations and Extensive Behavioral Profiling Tackling the Hippocampus Gradient. Cerebral Cortex, 29(11), 4595–4612. 10.1093/cercor/bhy336

Prompiengchai, S., & Dunlop, K. (2025). Breakthroughs and challenges for generating brain network-based biomarkers of treatment response in depression. Neuropsychopharmacology, 50(1), 230–245. 10.1038/s41386-024-01907-1

Raichle, M. E., MacLeod, A. M., Snyder, A. Z., Powers, W. J., Gusnard, D. A., & Shulman, G. L. (2001). A default mode of brain function. Proceedings of the National Academy of Sciences, 98(2), 676–682. 10.1073/pnas.98.2.676

Rajkowska, G., & Goldman-Rakic, P. S. (1995). Cytoarchitectonic Definition of Prefrontal Areas in the Normal Human Cortex: II. Variability in Locations of Areas 9 and 46 and Relationship to the Talairach Coordinate System. Cerebral Cortex, 5(4), 323–337. 10.1093/cercor/5.4.323

Reid, A. T., Bzdok, D., Langner, R., Fox, P. T., Laird, A. R., Amunts, K., Eickhoff, S. B., & Eickhoff, C. R. (2016). Multimodal connectivity mapping of the human left anterior and posterior lateral prefrontal cortex. Brain Structure and Function, 221(5), 2589–2605. 10.1007/s00429-015-1060-5

Reuter, N., Genon, S., Masouleh, S. K., Hoffstaedter, F., Liu, X., Kalenscher, T., Eickhoff, S. B., & Patil, K. R. (2020). CBPtools: a Python package for regional connectivity-based parcellation. Brain Structure and Function, 225(4), 1261–1275. 10.1007/s00429-020-02046-1

Sacchet, M. D., Ho, T. C., Connolly, C. G., Tymofiyeva, O., Lewinn, K. Z., Han, L. K., Blom, E. H., Tapert, S. F., Max, J. E., Frank, G. K., Paulus, M. P., Simmons, A. N., Gotlib, I. H., & Yang, T. T. (2016). Large-Scale Hypoconnectivity Between Resting-State Functional Networks in Unmedicated Adolescent Major Depressive Disorder. Neuropsychopharmacology, 41(12), 2951–2960. 10.1038/npp.2016.76

Sakreida, K., Trapp, N. T., Kreuzer, S., Rubin, U., Schnabel, D., Hovančáková, J., Sack, A. T., Neuner, I., Frodl, T., & Poeppl, T. B. (2025). Comparison of effectiveness of common targeting heuristics in repetitive transcranial magnetic stimulation treatment of depression. BMJ Mental Health, 28(1), e301598. 10.1136/bmjment-2025-301598

Salimi-Khorshidi, G., Douaud, G., Beckmann, C. F., Glasser, M. F., Griffanti, L., & Smith, S. M. (2014). Automatic denoising of functional MRI data: Combining independent component analysis and hierarchical fusion of classifiers. NeuroImage, 90, 449–468. 10.1016/j.neuroimage.2013.11.046

Sallet, J., Mars, R. B., Noonan, M. P., Neubert, F.-X., Jbabdi, S., O’Reilly, J. X., Filippini, N., Thomas, A. G., & Rushworth, M. F. (2013). The Organization of Dorsal Frontal Cortex in Humans and Macaques. The Journal of Neuroscience, 33(30), 12255–12274. 10.1523/jneurosci.5108-12.2013

Sarkissov, S. A., Filimonoff, I. N., Kononowa, E. P., Preobraschenskaja, I. S., & Kukuew, L. A. (1955). Atlas of Cytoarchitectonics of the Adult Human Cerebral Cortex. Medgiz.

Scalabrini, A., Vai, B., Poletti, S., Damiani, S., Mucci, C., Colombo, C., Zanardi, R., Benedetti, F., & Northoff, G. (2020). All roads lead to the default-mode network—global source of DMN abnormalities in major depressive disorder. Neuropsychopharmacology, 45(12), 2058–2069. 10.1038/s41386-020-0785-x

Seeley, W. W., Menon, V., Schatzberg, A. F., Keller, J., Glover, G. H., Kenna, H., Reiss, A. L., & Greicius, M. D. (2007). Dissociable Intrinsic Connectivity Networks for Salience Processing and Executive Control. The Journal of Neuroscience, 27(9), 2349–2356. 10.1523/jneurosci.5587-06.2007

Sheline, Y. I., Barch, D. M., Price, J. L., Rundle, M. M., Vaishnavi, S. N., Snyder, A. Z., Mintun, M. A., Wang, S., Coalson, R. S., & Raichle, M. E. (2009). The default mode network and self-referential processes in depression. Proceedings of the National Academy of Sciences, 106(6), 1942–1947. 10.1073/pnas.0812686106

Siddiqi, S. H., Schaper, F. L. W. V. J., Horn, A., Hsu, J., Padmanabhan, J. L., Brodtmann, A., Cash, R. F. H., Corbetta, M., Choi, K. S., Dougherty, D. D., Egorova, N., Fitzgerald, P. B., George, M. S., Gozzi, S. A., Irmen, F., Kuhn, A. A., Johnson, K. A., Naidech, A. M., Pascual-Leone, A.,…Fox, M. D. (2021). Brain stimulation and brain lesions converge on common causal circuits in neuropsychiatric disease. Nature Human Behaviour, 5(12), 1707–1716. 10.1038/s41562-021-01161-1

Siddiqi, S. H., Taylor, S. F., Cooke, D., Pascual-Leone, A., George, M. S., & Fox, M. D. (2020). Distinct Symptom-Specific Treatment Targets for Circuit-Based Neuromodulation. American Journal of Psychiatry, 177(5), 435–446. 10.1176/appi.ajp.2019.19090915

Smith, S. M., Beckmann, C. F., Andersson, J., Auerbach, E. J., Bijsterbosch, J., Douaud, G., Duff, E., Feinberg, D. A., Griffanti, L., Harms, M. P., Kelly, M., Laumann, T., Miller, K. L., Moeller, S., Petersen, S., Power, J., Salimi-Khorshidi, G., Snyder, A. Z., Vu, A. T.,…for the WU-Minn HCP Consortium. (2013). Resting-state fMRI in the Human Connectome Project. NeuroImage, 80, 144–168. 10.1016/j.neuroimage.2013.05.039

Sridharan, D., Levitin, D. J., & Menon, V. (2008). A critical role for the right fronto-insular cortex in switching between central-executive and default-mode networks. Proceedings of the National Academy of Sciences, 105(34), 12569–12574. 10.1073/pnas.0800005105

Thielscher, A., Antunes, A., & Saturnino, G. B. (2015). Field Modeling for Transcranial Magnetic Stimulation: A Useful Tool to Understand the Physiological Effects of TMS?**All authors contributed equally to the work. 2015 37th Annual International Conference of the IEEE Engineering in Medicine and Biology Society (EMBC), 2015, 222–225. 10.1109/embc.2015.7318340

Tripathi, V., Batta, I., Zamani, A., Atad, D. A., Sheth, S. K. S., Zhang, J., Wager, T. D., Whitfield-Gabrieli, S., Uddin, L. Q., Prakash, R. S., & Bauer, C. C. C. (2025). Default Mode Network Functional Connectivity As a Transdiagnostic Biomarker of Cognitive Function. Biological Psychiatry: Cognitive Neuroscience and Neuroimaging, 10(4), 359–368. 10.1016/j.bpsc.2024.12.016

Uddin, L. Q. (2015). Salience processing and insular cortical function and dysfunction. Nature Reviews Neuroscience, 16(1), 55–61. 10.1038/nrn3857

Uddin, L. Q., Yeo, B. T. T., & Spreng, R. N. (2019). Towards a Universal Taxonomy of Macro-scale Functional Human Brain Networks. Brain Topography, 32(6), 926–942. 10.1007/s10548-019-00744-6

U.S. Food and Drug Administration. (2018). *Major Depressive Disorder: Developing Drugs for Treatment*. U.S. Department of Health and Human Services Food and Drug Administration Center for Drug Evaluation and Research. https://www.fda.gov/media/113988/download

Van Essen, D. C., Smith, S. M., Barch, D. M., Behrens, T. E. J., Yacoub, E., Ugurbil, K., & for the WU-Minn HCP Consortium. (2013). The WU-Minn Human Connectome Project: An overview. NeuroImage, 80, 62–79. 10.1016/j.neuroimage.2013.05.041

Vinh, N. X., Epps, J., & Bailey, J. (2010). Information Theoretic Measures for Clusterings Comparison: Variants, Properties, Normalization and Correction for Chance. Journal of Machine Learning Research, 11(95), 2837–2854. https://jmlr.org/papers/v11/vinh10a.html

Von Economo, C. F., & Koskinas, G. N. (1925). Die Cytoarchitektonik der Hirnrinde des erwachsenen Menschen. Springer.

Vos de Wael, R., Benkarim, O., Paquola, C., Lariviere, S., Royer, J., Tavakol, S., Xu, T., Hong, S.-J., Langs, G., Valk, S., Misic, B., Milham, M., Margulies, D., Smallwood, J., & Bernhardt, B. C. (2020). BrainSpace: a toolbox for the analysis of macroscale gradients in neuroimaging and connectomics datasets. Communications Biology, 3(1), 103. 10.1038/s42003-020-0794-7

Weigand, A., Horn, A., Caballero, R., Cooke, D., Stern, A. P., Taylor, S. F., Press, D., Pascual-Leone, A., & Fox, M. D. (2018). Prospective Validation That Subgenual Connectivity Predicts Antidepressant Efficacy of Transcranial Magnetic Stimulation Sites. Biological Psychiatry, 84(1), 28–37. 10.1016/j.biopsych.2017.10.028

World Health Organization. (2017). Depression and Other Common Mental Disorders. World Health Organization.

Yan, C.-G., Chen, X., Li, L., Castellanos, F. X., Bai, T.-J., Bo, Q.-J., Cao, J., Chen, G.-M., Chen, N.-X., Chen, W., Cheng, C., Cheng, Y.-Q., Cui, X.-L., Duan, J., Fang, Y.-R., Gong, Q.-Y., Guo, W.-B., Hou, Z.-H., Hu, L.,…Zang, Y.-F. (2019). Reduced default mode network functional connectivity in patients with recurrent major depressive disorder. Proceedings of the National Academy of Sciences, 116(18), 9078–9083. 10.1073/pnas.1900390116

Yang, H., Chen, X., Chen, Z.-B., Li, L., Li, X.-Y., Castellanos, F. X., Bai, T.-J., Bo, Q.-J., Cao, J., Chang, Z.-K., Chen, G. M., Chen, N.-X., Chen, W., Cheng, C., Cheng, Y.-Q., Cui, X.-L., Duan, J., Fang, Y., Gong, Q.-Y.,…Yan, C.-G. (2021). Disrupted intrinsic functional brain topology in patients with major depressive disorder. Molecular Psychiatry, 26(12), 7363–7371. 10.1038/s41380-021-01247-2

Yeo, B. T. T., Krienen, F. M., Sepulcre, J., Sabuncu, M. R., Lashkari, D., Hollinshead, M., Roffman, J. L., Smoller, J. W., Zöllei, L., Polimeni, J. R., Fischl, B., Liu, H., & Buckner, R. L. (2011). The organization of the human cerebral cortex estimated by intrinsic functional connectivity. Journal of Neurophysiology, 106(3), 1125–1165. 10.1152/jn.00338.2011

Yu, M., Linn, K. A., Shinohara, R. T., Oathes, D. J., Cook, P. A., Duprat, R., Moore, T. M., Oquendo, M. A., Phillips, M. L., McInnis, M., Fava, M., Trivedi, M. H., McGrath, P., Parsey, R., Weissman, M. M., & Sheline, Y. I. (2019). Childhood trauma history is linked to abnormal brain connectivity in major depression. Proceedings of the National Academy of Sciences, 116(17), 8582–8590. 10.1073/pnas.1900801116

