## Supplementary Material for "Unveiling the Multifaceted Networks of the TMS-Targeted Left Prefrontal Cortex for Precision Neurostimulation"

### Supplementary Methods

**Table S1. Transcranial magnetic stimulation (TMS) coordinates used to create our region of interest**

| Site | MNIx | MNIy | MNIz | Reference |
|---|---|---|---|---|
| 5-cm average | -41 | 16 | 54 | (Fox et al., 2012) |
| 5.5-cm average | -33 | 30 | 50 | (Weigand et al., 2018) |
| Beam F3 average | -43 | 46 | 32 | (Cash et al., 2019) |
| 5-cm-Method–derived non-responders (posterior-medial position) | -41 | 17 | 55 | (Herbsman et al., 2009) |
| 5-cm-Method–derived responders (anterolateral position) | -46 | 23 | 49 | (Herbsman et al., 2009) |
| EEG F3 TMS target | -37 | 26 | 49 | (Herwig et al., 2003) |
| BA9/46 junction TMS target #1 | -46 | 45 | 38 | (Fitzgerald et al., 2009) |
| BA9/46 junction TMS target #2 | -50 | 30 | 36 | (Rusjan et al., 2010) |
| BA9 definition | -36 | 39 | 43 | (Rajkowska & Goldman-Rakic, 1995) |
| BA46 definition | -44 | 40 | 29 | (Rajkowska & Goldman-Rakic, 1995) |
| SGC-FC–derived group target #1 | -44 | 38 | 34 | (Fox et al., 2012) |
| SGC-FC–derived group target #2 | -38 | 44 | 26 | (Fox et al., 2012) |
| SGC-FC–derived group target #1 (N=1000) | -42 | 44 | 30 | (Weigand et al., 2018) |
| SGC-FC–derived group target (HCP dataset, N=2000) | -41 | 43 | 27 | (Cash, Cocchi, et al., 2021) |
| TMS Target | -40 | 31 | 34 | (Cho & Strafella, 2009; Paus et al., 2001) |

Common transcranial magnetic stimulation (TMS) coordinates targeting the left dorsolateral prefrontal cortex to treat depression (Cash, Cocchi, et al., 2021; Cash, Weigand, et al., 2021; Fox et al., 2012), which were used to create our region of interest for regional connectivity-based parcellation. **Abbreviations.** BA: Brodmann area, EEG: electroencephalogram, FC: functional connectivity, HCP: human connectome project, SGC: subgenual cingulate cortex, TMS: transcranial magnetic stimulation.

### 1. Internal and External Validity Metrics

The **Silhouette index** (Rousseeuw, 1987) is computed as follows: First, the mean distance between each pair of voxels within the same cluster (i.e., mean intra-cluster distance) is calculated. Second, the smallest mean distance between each voxel and all voxels in a different cluster (i.e., mean nearest-cluster distance) is identified. The Silhouette index is obtained by subtracting the mean intra-cluster distance from the mean nearest-cluster distance and dividing the result by the maximum value between both distances. The index ranges from -1 to 1, where higher values indicate better-defined clusters, i.e., higher separation between neighboring clusters (high inter-cluster distance) while keeping the dispersion within the cluster low (low intra-cluster distance). Values near 0 suggest overlapping clusters, whereas negative values indicate misclassified voxels, implying that a different number of clusters may better fit the data.

The **Calinski-Harabasz index**, or Variance Ratio Criterion, is calculated by, first, computing the intra-cluster distance as the sum of squared distances between each voxel and its cluster centroid. Next, the overall data centroid is computed and the inter-cluster distance is calculated as the sum of squared differences between the cluster centroids and the overall centroid. The inter-cluster distance is multiplied by the number of voxels minus the number of clusters. In turn, the intra-cluster distance is multiplied by the number of clusters minus 1. The Calinski-Harabasz index is finally obtained by dividing these two elements. Higher values indicate better clustering quality, with well-separated (high inter-cluster distance) but dense clusters (low intra-cluster distance).

The **Davies-Bouldin index** is calculated by first computing the mean distance between each voxel and the centroid of its cluster (i.e., intra-cluster distance). Then, the distance between each pair of cluster centroids (i.e., inter-cluster distance) is computed. For each pair of clusters, the ratio between the sum of their intra-cluster distances and the distance between their centroids is computed. The maximal ratio is selected for each cluster, and the Davies-Bouldin index is the average of all maximal ratio values across all clusters. As such, this metric has a minimum value of 0 without an upper bound, where lower values indicate better clustering by having high compactness (low intra-cluster distance) and high separation (high inter-cluster distance).

The **Adjusted Rand Index (ARI)** measures clustering similarity while accounting for random overlaps, comparing the number of voxel pairs assigned to the same or different clusters relative to what would be expected under a random assignment. ARI is a symmetric score ranging from -1 to 1, where higher values indicate greater alignment between individual- and group-level clustering solutions. Values near 0 suggest that clustering similarity is no better than random assignment, whereas negative values indicate that the clustering structure is worse than expected by chance.

The **Adjusted Mutual Information (AMI)** quantifies how much information is shared between two clustering solutions, i.e., the reduction in uncertainty about one clustering given knowledge of another clustering (Vinh et al., 2010). The correction accounts for chance and the tendency of raw mutual information to increase with the number of clusters, regardless of whether more information is actually shared (Eickhoff et al., 2015). AMI takes values between 0 and 1, where 0 denotes clustering solutions that agree no better than chance, and 1 denotes identical clusterings and maximal reduction of uncertainty. Being information-theoretic, AMI is invariant to arbitrary cluster labeling, distinguishing it from the pair-counting, index-based ARI.

The **Normalized Variation of Information (NVI)** quantifies the information-theoretic distance between two clustering solutions. It is defined as the sum of the conditional entropies in both directions, i.e., the information about one solution that is not conveyed by the other, or the leftover uncertainty in both directions (Meilă, 2007). Opposite to the raw mutual information, the raw variation of information favors solutions with lower *k*. We therefore normalized it by the joint entropy, yielding values bounded between 0 and 1 that express the leftover uncertainty relative to the total information present in both partitions (Kraskov et al., 2005). Lower values indicate greater correspondence, with 0 denoting identical partitions. Importantly, the normalization does not correct for chance, and, unlike AMI, NVI is expected to drift with *k* under random labeling. Therefore, it is interpreted based on the trend across *k* rather than in absolute terms. Like AMI, NVI is independent of cluster labeling.

The **relabeling accuracy** is the ratio of label overlaps between the individual and reference clusterings.

#### 2. Difference Likelihood Map of TMS-PFC According to SGC Connectivity

As a complementary approach to the overlap map (minimal conjunction) reported in the main text (Section 3.4, **Figure 6A**), we quantified each voxel's likelihood of showing positive or negative SGC connectivity in a single continuous map by generating a difference likelihood map. Whereas the conjunction yields a binary delineation of voxels assigned to the respective cluster at every granularity, this probability-weighted approach additionally incorporates between-subject consistency of cluster assignment, thereby representing the antagonistic organization as a continuous spatial gradient across the ROI. The resulting difference likelihood map thus shows the relative likelihood of TMS-PFC areas showing positive or negative SGC connectivity across granularities and individuals, denoted SGC+ and SGC-.

To obtain this map, the respective between-subject cluster probability maps (see *Methods—Regional Connectivity-Based Parcellation* and **Supplementary Figure S3** as an example for $k$=9) were averaged across granularities to obtain two composite maps representing the likelihood of belonging to the TMS-PFC regions most positively (SGC+) or negatively (SGC-) connected with SGC. We then subtracted the SGC- map from the SGC+ map across the entire ROI, yielding the difference likelihood map. Positive values indicate stronger consistency of SGC+ connectivity, while negative values indicate stronger consistency of SGC- connectivity across granularities and individuals (**Figure 6B** and **Supplementary Figure S6A**). These SGC+ and SGC- areas resemble the locations of clusters one and six in the nine-cluster solution, but appear slightly shifted and spatially more extended as a result of merging across granularities. To isolate the most spatially reliable regions across individuals, we thresholded the difference map at ±50% (delineated in white in **Supplementary Figure S6A** and shown in red and blue at the top of **Supplementary Figure S6B**). Through this thresholding, the resulting extreme areas are somewhat smaller than those obtained by the overlap map (**Figure 6A**). These extreme SGC+ and SGC- areas were then analyzed in terms of their whole-brain functional connectivity and behavioral associations, as reported in Supplementary Results (**Supplementary Figure S6**).

#### 2. Assessment of Spatial Coherence of Cluster Solutions

To assess the spatial continuity of each clustering solution, we identified spatially disconnected components within each cluster label using a 26-connectivity criterion (i.e., voxels sharing a face, edge, or corner with at least one other voxel of the same label were considered part of the same component).

For each granularity level ($k$ = 2–10) and each cluster within that granularity, we counted the number of connected components per cluster label and their corresponding number of voxels. A clustering solution was considered spatially discontinuous if any of its clusters contained more than one continuous component. A secondary component, that is, a spatially separate component disconnected from the main cluster body, was defined as one exceeding five voxels. This threshold was chosen to tolerate negligible border artifacts arising from the parcellation procedure while flagging meaningful spatial discontinuities. Granularities containing at least one such discontinuous cluster were excluded from subsequent SGC connectivity analyses.

### Supplementary Results

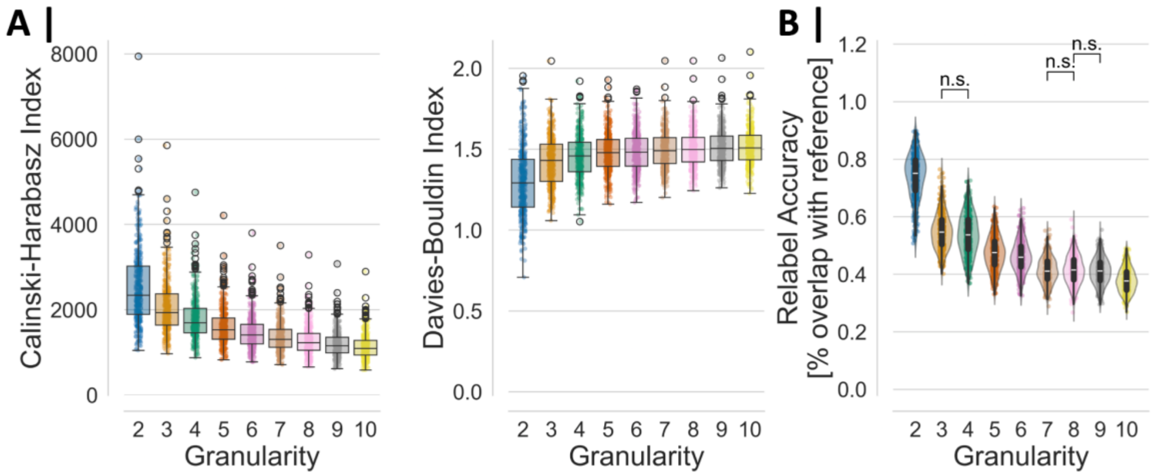


**Figure S1. Internal and external validity metrics for regional connectivity-based parcellation of the TMS-targeted left prefrontal cortex. A.** Internal validity scores for each granularity level. Higher Calinski-Harabasz index values indicate greater separation and compactness of clusters, reflecting higher similarity between the individual source RSFC data and the group-level reference clustering image. In contrast, lower Davies-Bouldin index values denote better clustering performance, with smaller intra-cluster variance relative to inter-cluster separation. **B.** External validity assessed via relabeling accuracy between the individual-level clustering after relabeling and the group-level reference clustering, indicating the consistency of the clustering across subjects. **Abbreviations.** n.s.: not significant ($p > 0.05$), TMS: transcranial magnetic stimulation.

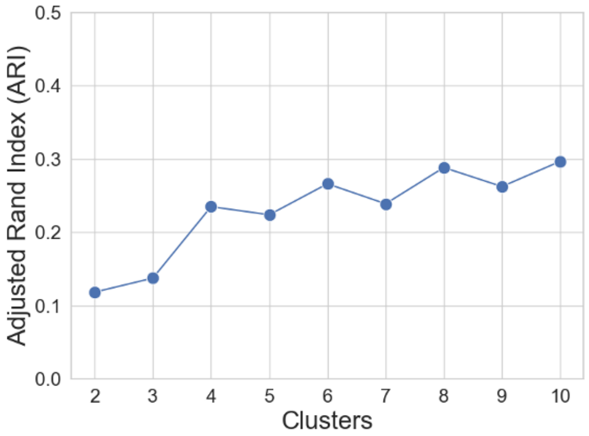


**Figure S2. Similarity between regional connectivity-based parcellation of the TMS-targeted left prefrontal cortex and the left DLPFC from the Jülich-Brain Cytoarchitectonic Atlas.** The Jülich-Brain Cytoarchitectonic Atlas was converted into MNI152NLin6Asym space with 2 mm isotropic voxels. Only overlapping voxels between the TMS-PFC seed and the cytoarchitectonic atlas were kept (3233 voxels), and the adjusted rand index (ARI) was computed between each parcellation result and the overlapping cytoarchitectonic regions. **Abbreviations.** DLPFC: dorsolateral prefrontal cortex, TMS: transcranial magnetic stimulation, TMS-PFC: TMS-targeted left prefrontal cortex.

**Table S2. Overlap between functional TMS-PFC subregions and cytoarchitectonic areas**

| *k* | cluster_id | ARI | cyto_name | overlap_voxels | overlap_ratio |
|---|---|---|---|---|---|
| 2 | **1** | 0.1186 | **Area 8v1 (MFG)** | **370** | **21.30** |
| | **2** | | **Area MFG4 (MFG)** | **554** | **37.03** |
| 3 | **1** | 0.1377 | **Area 8v1 (MFG)** | **362** | **24.95** |
| | 2 | | Area SFG4 (SFG) | 141 | 22.49 |
| | **3** | | **Area MFG4 (MFG)** | **419** | **36.28** |
| 4 | 1 | 0.2352 | Area 8v1 (MFG) | 359 | 32.94 |
| | 2 | | Area 8v2 (MFG) | 107 | 30.31 |
| | 3 | | Area MFG4 (MFG) | 505 | 51.53 |
| | 4 | | Area 45 (IFG) | 191 | 23.58 |
| 5 | 1 | 0.2241 | Area 8v1 (MFG) | 326 | 31.62 |
| | 2 | | Area 8v1 (MFG) | 51 | 40.8 |
| | 3 | | Area SFG4 (SFG) | 139 | 27.58 |
| | 4 | | Area MFG4 (MFG) | 446 | 58.84 |
| | 5 | | Area 45 (IFG) | 194 | 23.80 |
| 6 | **1** | 0.2664 | **Area 8d2 (SFG)** | **239** | **34.39** |
| | 2 | | Area SFG4 (SFG) | 142 | 31.98 |
| | **3** | | **Area MFG4 (MFG)** | **364** | **57.59** |
| | 4 | | Area 8v1 (MFG) | 203 | 39.65 |
| | 5 | | Area MFG5 (MFG, IFG) | 178 | 39.73 |
| | 6 | | Area 45 (IFG) | 158 | 31.47 |

| | | | | | |
|---|---|---|---|---|---|
| 7 | 1 | 0.2391 | Area 8v1 (MFG) | 242 | 30.36 |
| | 2 | | Area SFG4 (SFG) | 168 | 76.02 |
| | 3 | | Frontal-II (GapMap) | 109 | 23.09 |
| | 4 | | Area MFG4 (MFG) | 233 | 42.67 |
| | 5 | | Area 8v1 (MFG) | 142 | 35.50 |
| | 6 | | Area MFG4 (MFG) | 242 | 63.19 |
| | 7 | | Area MFG5 (MFG, IFG) | 161 | 38.89 |
| 8 | **1** | 0.2885 | **Area 8d2 (SFG)** | **239** | **36.32** |
| | 2 | | Area SFG4 (SFG) | 157 | 35.04 |
| | **3** | | **Area MFG4 (MFG)** | **342** | **60.32** |
| | 4 | | Area 8v1 (MFG) | 222 | 54.68 |
| | 5 | | Frontal-II (GapMap) | 104 | 40.31 |
| | 6 | | Area MFG5 (MFG, IFG) | 132 | 47.14 |
| | 7 | | Area 45 (IFG) | 119 | 30.05 |
| | 8 | | Area 45 (IFG) | 75 | 34.09 |
| 9 | **1** | 0.2626 | **Area 8d2 (SFG)** | **234** | **38.11** |
| | 2 | | Area MFG4 (MFG) | 102 | 26.22 |
| | 3 | | Area SFG4 (SFG) | 110 | 93.22 |
| | 4 | | Area 8v2 (MFG) | 96 | 38.10 |
| | 5 | | Frontal-II (GapMap) | 90 | 36.59 |
| | **6** | | **Area MFG4 (MFG)** | **280** | **64.52** |
| | 7 | | Area 8v1 (MFG) | 166 | 47.16 |

| | | | | | |
|---|---|---|---|---|---|
| | 8 | | Area MFG5 (MFG, IFG) | 181 | 44.91 |
| | 9 | | Area 45 (IFG) | 127 | 29.88 |
| 10 | 1 | 0.297 | Area 8d2 (SFG) | 124 | 36.9 |
| | 2 | | Area 8v1 (MFG) | 121 | 36.01 |
| | 3 | | Area SFG4 (SFG) | 131 | 60.09 |
| | 4 | | Area MFG4 (MFG) | 121 | 31.11 |
| | 5 | | Area MFG4 (MFG) | 339 | 67.4 |
| | 6 | | Area 8v1 (MFG) | 191 | 60.06 |
| | 7 | | Area MFG5 (MFG, IFG) | 120 | 44.44 |
| | 8 | | Area MFG5 (MFG, IFG) | 71 | 33.18 |
| | 9 | | Frontal-II (GapMap) | 125 | 47.17 |
| | 10 | | Area 45 (IFG) | 120 | 31.25 |

**Note.** The overlap ratio was obtained by dividing the number of overlapped voxels by the total number of voxels in the functional cluster. Bold areas highlight clusters that showed the strongest positive and negative connectivity with the subgenual cingulate cortex in granularities without spatially discontinuous clusters ($k$ = 2, 3, 6, 8, 9). **Abbreviations.** IFG: inferior frontal gyrus, MFG: middle frontal gyrus, SFG: superior frontal gyrus, TMS: transcranial magnetic stimulation, TMS-PFC: TMS-targeted left prefrontal cortex.

133

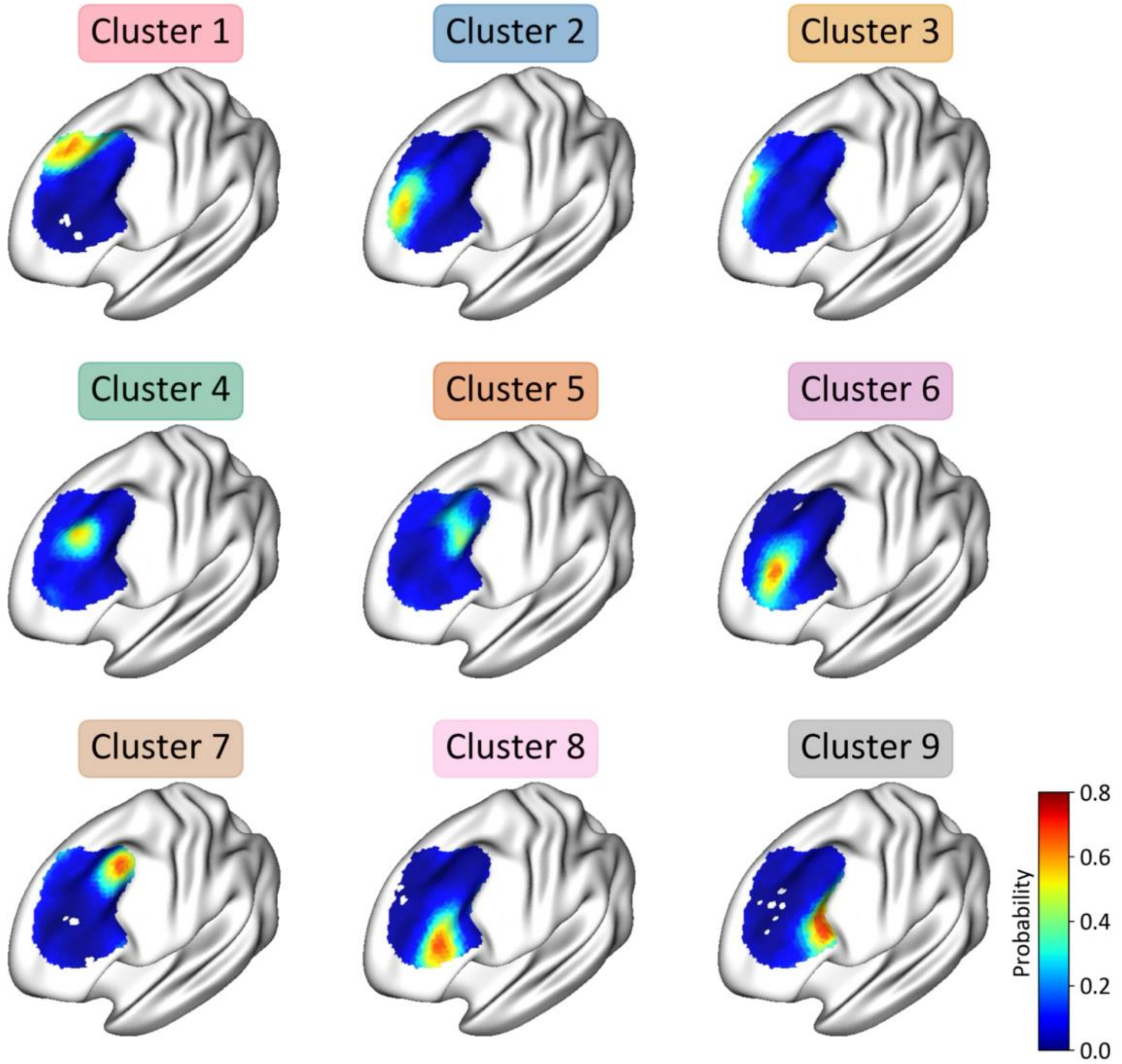


**Figure S3. Probabilistic map for each cluster in the nine-cluster solution of the TMS-targeted left prefrontal cortex.** Maps represent the voxel-wise likelihood of cluster assignment across all subjects after relabeling individual cluster labels with the reference clustering image. **Abbreviations.** TMS: transcranial magnetic stimulation.

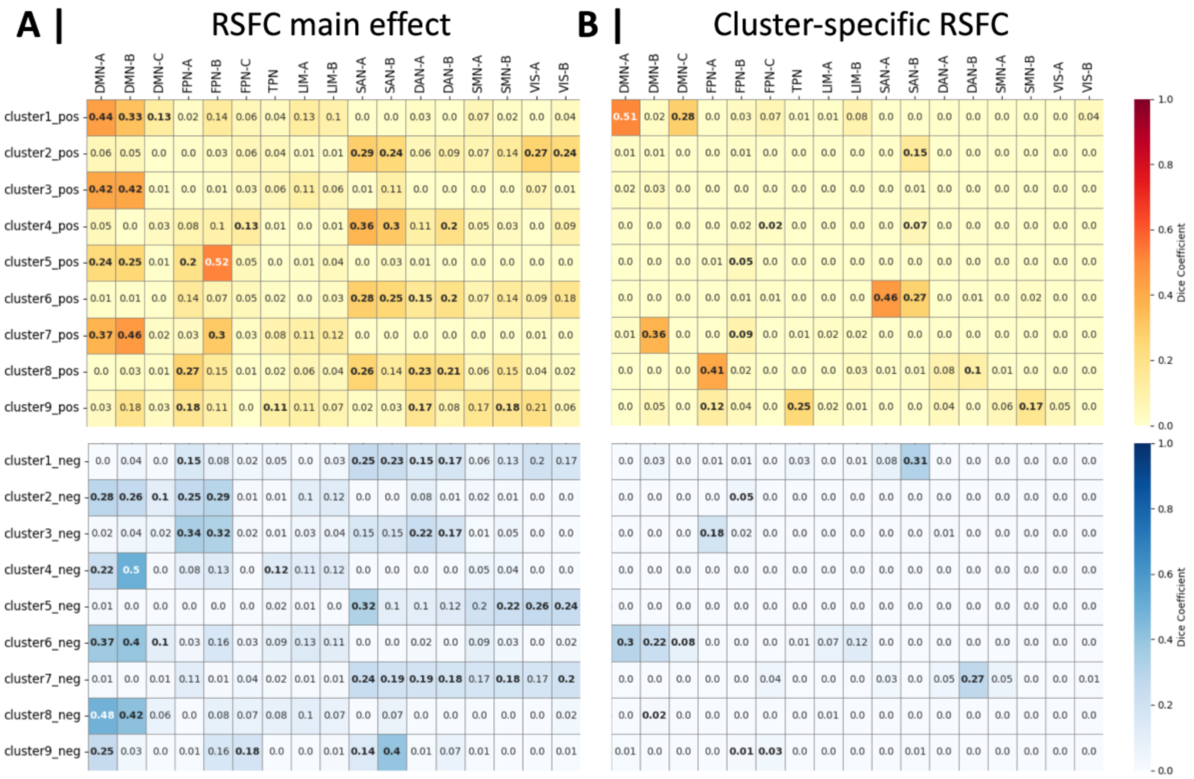


**Figure S4. Network correspondence of TMS-PFC subregions' RSFC profiles with 17-Yeo networks. A.** Correspondence between the positive (top) and negative (bottom) RSFC main effects for each cluster of the nine-cluster solution and the 17-Yeo networks. **B.** Correspondence between the positive (top) and negative (bottom) cluster-specific RSFC profiles for each cluster of the nine-cluster solutions and the 17-Yeo networks. Bold labels denote statistically significant overlap after spin-test permutations. **Abbreviations.** DAN: dorsal attention network, DMN: default-mode network, FPN: frontoparietal network, LIM: limbic network, neg: negative, pos: positive, RSFC: resting-state functional connectivity, SAN: salience attention network, SMN: somatomotor network, TMS: transcranial magnetic stimulation, TMS-PFC: TMS-targeted left prefrontal cortex, TPN: temporoparietal network, VIS: visual network.

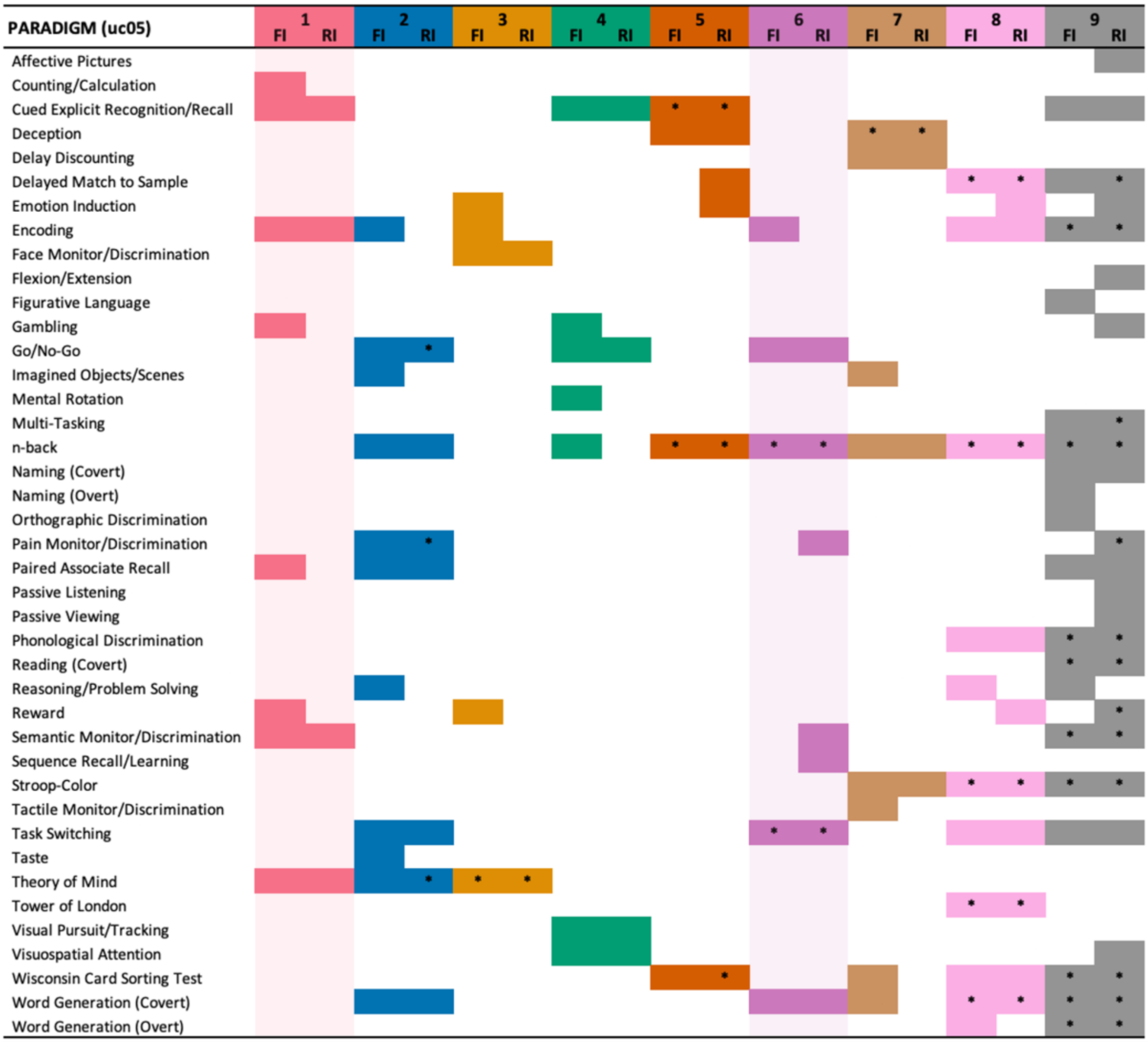


**Figure S5. Behavioral functional profiling of the TMS-targeted left prefrontal cortex subregions.** Paradigms associated with each cluster in the nine-cluster solution. The shaded sectors emphasize the clusters with strongest positive (cluster one) and negative (cluster six) connectivity with the subgenual cingulate cortex. **Abbreviations.** FI: forward inference as P(Activation|Task), RI: reverse inference as P(Task|Activation), TMS: transcranial magnetic stimulation, uc05: uncorrected *p* < 0.05, * FDR-corrected *p* < 0.05.

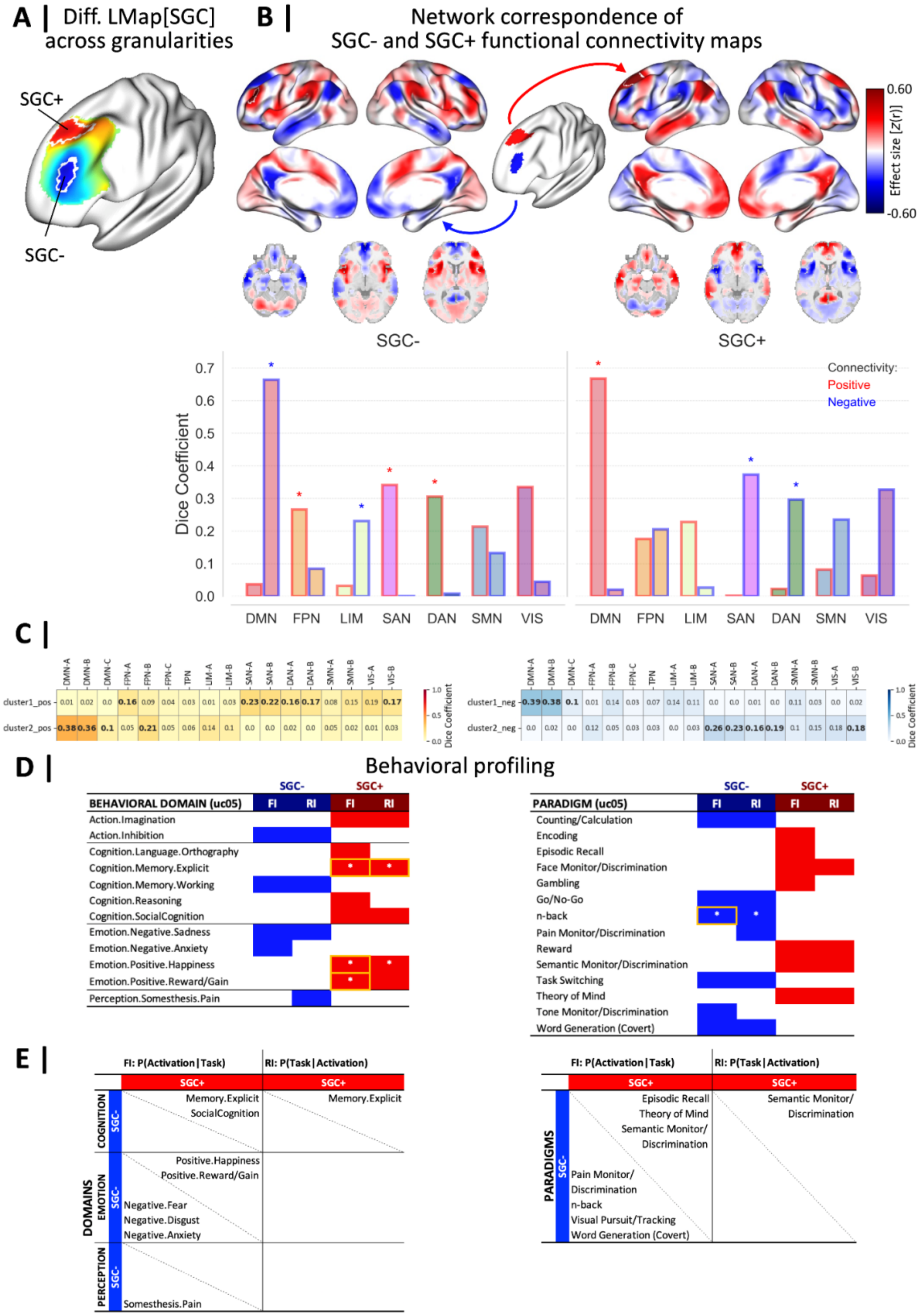

A | Diff. LMap[SGC] across granularities
SGC+
SGC-
B | Network correspondence of SGC- and SGC+ functional connectivity maps
0.60
Effect size [Z(r)]
-0.60
SGC-
SGC+
Dice Coefficient
Connectivity:
Positive
Negative
DMN FPN LIM SAN DAN SMN VIS
C |
DMN-A DMN-B DMN-C FPN-A FPN-B FPN-C TPN LIM-A LIM-B SAN-A SAN-B DAN-A DAN-B SMN-A SMN-B VIS-A VIS-B
cluster1_pos 0.01 0.02 0.0 0.16 0.09 0.04 0.03 0.01 0.03 0.23 0.22 0.16 0.17 0.08 0.15 0.19 0.17
cluster2_pos 0.38 0.36 0.1 0.05 0.21 0.05 0.06 0.14 0.1 0.0 0.0 0.02 0.0 0.05 0.02 0.01 0.03
cluster1_neg 0.39 0.38 0.1 0.01 0.14 0.03 0.07 0.14 0.11 0.0 0.0 0.01 0.0 0.11 0.03 0.0 0.02
cluster2_neg 0.0 0.02 0.0 0.12 0.05 0.03 0.03 0.0 0.03 0.26 0.23 0.16 0.19 0.1 0.15 0.18 0.18
Dice Coefficient
D | Behavioral profiling
SGC- SGC+
BEHAVIORAL DOMAIN (uc05) FI RI FI RI
Action.Imagination
Action.Inhibition
Cognition.Language.Orthography
Cognition.Memory.Explicit
Cognition.Memory.Working
Cognition.Reasoning
Cognition.SocialCognition
Emotion.Negative.Sadness
Emotion.Negative.Anxiety
Emotion.Positive.Happiness
Emotion.Positive.Reward/Gain
Perception.Somesthesis.Pain
PARADIGM (uc05) FI RI FI RI
Counting/Calculation
Encoding
Episodic Recall
Face Monitor/Discrimination
Gambling
Go/No-Go
n-back
Pain Monitor/Discrimination
Reward
Semantic Monitor/Discrimination
Task Switching
Theory of Mind
Tone Monitor/Discrimination
Word Generation (Covert)
E |
FI: P(Activation|Task) RI: P(Task|Activation)
SGC+
DOMAINS
COGNITION SGC-
Memory.Explicit
SocialCognition
Memory.Explicit
EMOTION SGC-
Positive.Happiness
Positive.Reward/Gain
Negative.Fear
Negative.Disgust
Negative.Anxiety
PERCEPTION SGC-
Somesthesis.Pain
PARADIGMS SGC-
Episodic Recall
Theory of Mind
Semantic Monitor/ Discrimination
Semantic Monitor/ Discrimination
Pain Monitor/ Discrimination
n-back
Visual Pursuit/Tracking
Word Generation (Covert)

**Figure S6. Functional characterization of difference likelihood map of TMS-PFC. A.** Difference likelihood map of TMS-PFC subregions according to their connectivity with SGC across granularities ($k$ = 2, 3, 6, 8, 9), obtained by averaging between-subject cluster probability maps (see **Supplementary Methods 2**). White borders delineate the extreme SGC- and SGC+ clusters (i.e., difference likelihood map thresholded at 50%). **B.** Positive and negative RSFC main effects for extreme SGC- and SGC+ clusters (i.e., thresholded at 50%) and their network correspondence (Dice coefficients) with the 7-Yeo networks. Bars with asterisks denote statistically significant overlap larger than expected by chance after spin-test permutations. **C.** Correspondence between the 17-Yeo networks and the positive (left) and negative (right) RSFC main effects for the extreme (1) SGC- and (2) SGC+ clusters. Bold annotations demarcate statistically significant overlap after spin-test permutations. **D.** Behavioral domains (left) and paradigm classes (right) associated with the extreme SGC- and SGC+ clusters shown in panel B. The orange frames mark the significant domains and paradigms (FDR-corrected) that also turned significant when analyzing the contrast between the two clusters. **E.** Contrast results of behavioral domains (left) and paradigm classes (right) from the behavioral functional profiling for the extreme SGC- and SGC+ clusters shown in panel B using BrainMap. Labels in the lower-left triangles indicate significantly stronger associations with SGC- (vs. SGC+), whereas those in the upper-right triangles indicate significantly stronger associations with SGC+ (vs. SGC-). Only significant results are shown (FDR-corrected $p < 0.05$). **Abbreviations.** DAN: dorsal attention network, DMN: default-mode network, FI: forward inference as P(Activation|Task), FPN: frontoparietal network, LIM: limbic network, neg: negative, pos: positive, RI: reverse inference as P(Task|Activation), RSFC: resting-state functional connectivity, SAN: salience attention network, SGC: subgenual cingulate cortex, TMS: transcranial magnetic stimulation, TMS-PFC: TMS-targeted left prefrontal cortex, uc05: uncorrected $p < 0.05$, * FDR-corrected $p < 0.05$. **Note.** SGC- is the TMS-PFC area with more than 50% likelihood of being anti-correlated with SGC across granularities, and SGC+ is the TMS-PFC area with more than 50% likelihood of having positive connectivity with SGC across granularities.

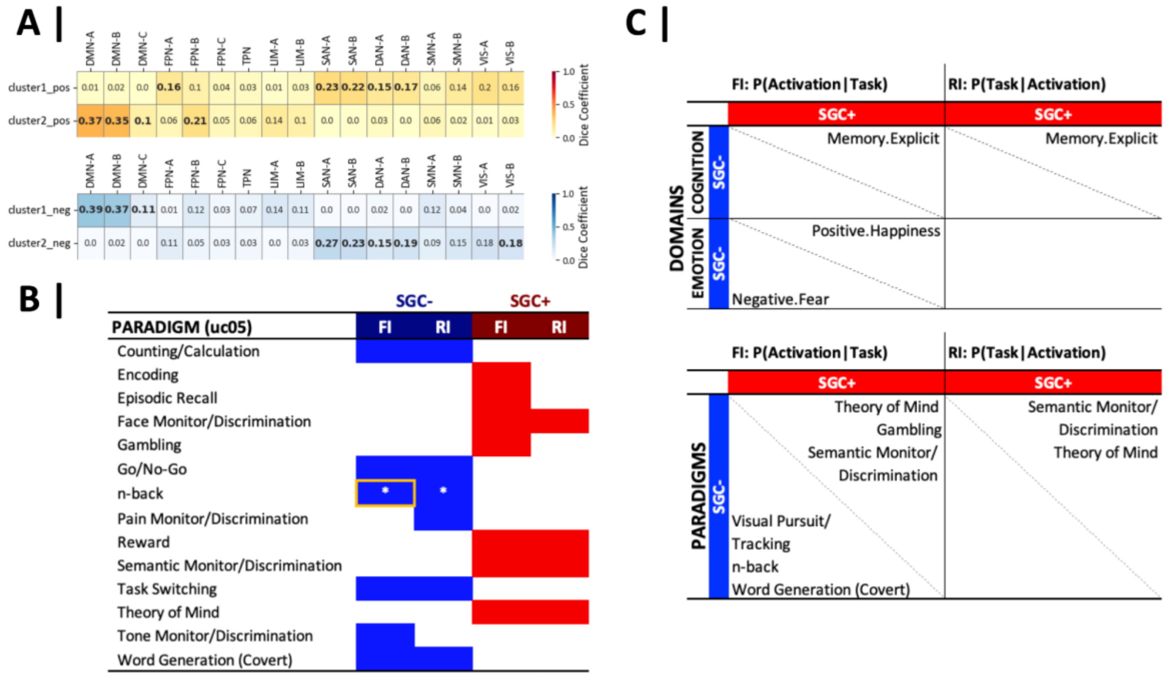


**Figure S7. Network correspondence with 17-Yeo networks and behavioral functional profiling based on paradigm classes and contrasts for TMS-PFC SGC- and SGC+ regions. A.** Correspondence between the 17-Yeo networks and the positive (top) and negative (bottom) RSFC main effects for the (1) SGC- and (2) SGC+ clusters shown in **Figure 6A**. Bold annotations demarcate statistically significant overlap after spin-test permutations. **B.** Paradigms associated with the SGC- and SGC+ clusters. The orange frame marks the significant paradigm (FDR-corrected) that also turned significant when analyzing the contrast between the two clusters. **C.** Contrast results of behavioral domains (left) and paradigm classes (right) from the behavioral functional profiling for the SGC- and SGC+ clusters using BrainMap. Labels in the lower-left triangles indicate significantly stronger associations with SGC- (vs. SGC+), whereas those in the upper-right triangles indicate significantly stronger associations with SGC+ (vs. SGC-). Only significant results are shown (FDR-corrected $p < 0.05$). **Abbreviations.** DAN: dorsal attention network, DMN: default-mode network, FI: forward inference as P(Activation|Task), FPN: frontoparietal network, LIM: limbic network, neg: negative, pos: positive, RI: reverse inference as P(Task|Activation), RSFC: resting-state functional connectivity, SAN: salience attention network, SGC: subgenual cingulate cortex, SMN: somatomotor network, TMS: transcranial magnetic stimulation, TMS-PFC: TMS-targeted left prefrontal cortex, TPN: temporoparietal network, uc05: uncorrected $p < 0.05$, VIS: visual network, * FDR-corrected $p < 0.05$.

#### References


Cash, R. F. H., Cocchi, L., Lv, J., Fitzgerald, P. B., & Zalesky, A. (2021). Functional Magnetic Resonance Imaging–Guided Personalization of Transcranial Magnetic Stimulation Treatment for Depression. *JAMA Psychiatry*, *78*(3), 337–339. https://doi.org/10.1001/jamapsychiatry.2020.3794

Cash, R. F. H., Weigand, A., Zalesky, A., Siddiqi, S. H., Downar, J., Fitzgerald, P. B., & Fox, M. D. (2021). Using Brain Imaging to Improve Spatial Targeting of Transcranial Magnetic Stimulation for Depression. *Biological Psychiatry*, *90*(10), 689–700. https://doi.org/10.1016/j.biopsych.2020.05.033

Cash, R. F. H., Zalesky, A., Thomson, R. H., Tian, Y., Cocchi, L., & Fitzgerald, P. B. (2019). Subgenual Functional Connectivity Predicts Antidepressant Treatment Response to Transcranial Magnetic Stimulation: Independent Validation and Evaluation of Personalization. *Biological Psychiatry*, *86*(2), e5–e7. https://doi.org/10.1016/j.biopsych.2018.12.002

Cho, S. S., & Strafella, A. P. (2009). rTMS of the Left Dorsolateral Prefrontal Cortex Modulates Dopamine Release in the Ipsilateral Anterior Cingulate Cortex and Orbitofrontal Cortex. *PLoS ONE*, *4*(8), e6725. https://doi.org/10.1371/journal.pone.0006725

Eickhoff, S. B., Thirion, B., Varoquaux, G., & Bzdok, D. (2015). Connectivity-based parcellation: Critique and implications. *Human Brain Mapping*, *36*(12), 4771–4792. https://doi.org/10.1002/hbm.22933

Fitzgerald, P. B., Hoy, K., McQueen, S., Maller, J. J., Herring, S., Segrave, R., Bailey, M., Been, G., Kulkarni, J., & Daskalakis, Z. J. (2009). A Randomized Trial of rTMS Targeted with MRI Based Neuro-Navigation in Treatment-Resistant Depression. *Neuropsychopharmacology*, *34*(5), 1255–1262. https://doi.org/10.1038/npp.2008.233

Fox, M. D., Buckner, R. L., White, M. P., Greicius, M. D., & Pascual-Leone, A. (2012). Efficacy of Transcranial Magnetic Stimulation Targets for Depression Is Related to Intrinsic Functional Connectivity with the Subgenual Cingulate. *Biological Psychiatry*, *72*(7), 595–603. https://doi.org/10.1016/j.biopsych.2012.04.028

Herbsman, T., Avery, D., Ramsey, D., Holtzheimer, P., Wadjik, C., Hardaway, F., Haynor, D., George, M. S., & Nahas, Z. (2009). More Lateral and Anterior Prefrontal Coil Location Is Associated with Better Repetitive Transcranial Magnetic Stimulation Antidepressant Response. *Biological Psychiatry*, *66*(5), 509–515. https://doi.org/10.1016/j.biopsych.2009.04.034

Herwig, U., Lampe, Y., Juengling, F. D., Wunderlich, A., Walter, H., Spitzer, M., & Schönfeldt-Lecuona, C. (2003). Add-on rTMS for treatment of depression: a pilot study using stereotaxic coil-navigation according to PET data. *Journal of Psychiatric Research*, *37*(4), 267–275. https://doi.org/10.1016/s0022-3956(03)00042-6

Kraskov, A., Stögbauer, H., Andrzejak, R. G., & Grassberger, P. (2005). Hierarchical clustering using mutual information. *EPL (Europhysics Letters)*, *70*(2), 278–284. https://doi.org/10.1209/epl/i2004-10483-y

Meilă, M. (2007). Comparing clusterings—an information based distance. *Journal of Multivariate Analysis*, *98*(5), 873–895. https://doi.org/10.1016/j.jmva.2006.11.013

Paus, T., Castro-Alamancos, M. A., & Petrides, M. (2001). Cortico-cortical connectivity of the human mid-dorsolateral frontal cortex and its modulation by repetitive transcranial magnetic stimulation. *European Journal of Neuroscience*, *14*(8), 1405–1411. https://doi.org/10.1046/j.0953-816x.2001.01757.x

Rajkowska, G., & Goldman-Rakic, P. S. (1995). Cytoarchitectonic Definition of Prefrontal Areas in the Normal Human Cortex: II. Variability in Locations of Areas 9 and 46 and Relationship to the Talairach Coordinate System. *Cerebral Cortex*, *5*(4), 323–337. https://doi.org/10.1093/cercor/5.4.323

Rousseeuw, P. J. (1987). Silhouettes: A graphical aid to the interpretation and validation of cluster analysis. *Journal of Computational and Applied Mathematics*, *20*, 53–65. https://doi.org/10.1016/0377-

0427(87)90125-7

Rusjan, P. M., Barr, M. S., Farzan, F., Arenovich, T., Maller, J. J., Fitzgerald, P. B., & Daskalakis, Z. J. (2010). Optimal transcranial magnetic stimulation coil placement for targeting the dorsolateral prefrontal cortex using novel magnetic resonance image-guided neuronavigation. *Human Brain Mapping*, *31*(11), 1643–1652. https://doi.org/10.1002/hbm.20964

Vinh, N. X., Epps, J., & Bailey, J. (2010). Information Theoretic Measures for Clusterings Comparison: Variants, Properties, Normalization and Correction for Chance. *Journal of Machine Learning Research*, *11*(95), 2837–2854. https://jmlr.org/papers/v11/vinh10a.html

Weigand, A., Horn, A., Caballero, R., Cooke, D., Stern, A. P., Taylor, S. F., Press, D., Pascual-Leone, A., & Fox, M. D. (2018). Prospective Validation That Subgenual Connectivity Predicts Antidepressant Efficacy of Transcranial Magnetic Stimulation Sites. *Biological Psychiatry*, *84*(1), 28–37. https://doi.org/10.1016/j.biopsych.2017.10.028
